# Horizontal gene transfers and terpene metabolism drive plant-fungal interaction in *Marchantia polymorpha*

**DOI:** 10.1101/2024.12.20.629586

**Authors:** Karima El Mahboubi, Chloé Beaulieu, Baptiste Castel, Cyril Libourel, Nathanaël Jariais, Emilie Amblard, Fabian van Beveren, Jean Keller, Yves Martinez, Jessica Nelson, Maxime Bonhomme, Christophe Jacquet, Pierre-Marc Delaux

**Affiliations:** Laboratoire de Recherche en Sciences Végétales (LRSV), Université de Toulouse, CNRS, UPS, Toulouse INP, Castanet-Tolosan, France; TRI-FRAIB Imaging Platform Facilities, FRAIB, Université de Toulouse, CNRS, UPS, 31320 Castanet-Tolosan, France; Natural History Museum, University of Oslo, Oslo, Norway

## Abstract

The liverwort *Marchantia polymorpha* has emerged as a model for studying plant immunity in bryophytes, providing unique insights into conserved defense mechanisms across land plants. By contrast, Marchantia-specific immune mechanisms have not been explored. In this study, we investigated the genetic basis of quantitative resistance in *M. polymorpha* against the fungal pathogen *Colletotrichum nymphaeae*, a naturally occurring compatible parasite. Through a combination of phenotypic, cytological and transcriptomic approaches, combined with genome-wide association studies (GWAS), we identified key defense-related genes and pathways. Transcriptomic analyses performed on two lines with contrasting suceptibilities to the pathogen revealed a strong overlap in immune responses between *M. polymorpha* and angiosperms, including the upregulation of PR proteins, transcription factors typically associated with biotic stresses, and enzymes involved in specialized metabolism. Leveraging the biological and genetic variability present in a collection of natural *M. polymorpha* accessions, highlight the role of horizontally transferred microbial-like terpene synthase (MTPSL) genes, which may contribute to the exceptional terpene diversity in liverworts and play a role in pathogen resistance. GWAS uncovered candidate loci associated with resistance traits, implicating both core immune components and specialized metabolic pathways. These results provide new insights into the specific molecular underpinnings of bryophyte immunity and underscore the evolutionary significance of horizontal gene transfer and specialized metabolites in shaping plant-pathogen interactions.

## Introduction

Half a billion years ago, plants transitioned from an aquatic to a terrestrial environment (Beerling., 2007). This habitat shift required the evolution of adaptations to abiotic challenges such as nutrient scarcity, or UV stress and drought (Rensing 2018). Simultaneously, biotic pressures from emerging pathogens drove the evolution and diversification of plant immune responses (Delaux & Schornack, 2021).

Extant land plant species are divided into two main lineages, the tracheophytes (vascular plants, which includes flowering plants), and the bryophytes that include liverworts, mosses and hornworts (Morris et al., 2018). These two lineages diverged from each other early after the colonization of lands (Morris et al., 2018), making possible to reconstruct the biology of the first land plants by comparing extant species belonging to the tracheophytes and bryophytes to identify similarities and homologies (Delaux et al., 2019). This allowed for instance the identification of an ancestral biosynthesis pathway for the plant cuticle (Knosp et al., 2024), the demonstration of the ancestral nature of symbiotic interactions (Rich et al., 2021) or the occurrence of a shared program for the development of epidermal structures across land plants (Proust et al., 2016).

While physiological and molecular mechanisms involved in plant immunity have been extensively studied in flowering plants (Ngou et al., 2022) our understanding of defense mechanisms in bryophytes remains limited. Phylogenetic surveys across the diversity of land plants (Han, 2019) and targeted functional analyses in model bryophytes have revealed the conservation of several immune components, including the involvement of Lysin-motif Receptor-like Kinase (LysM-RLK) receptors for the perception of fungal pathogens and the activation of downstream MAP-Kinases (Bressendorf et al., 2016, Yotsui et al., 2023), NPR1 and salicylic acid (SA) as key elements in hormonal immune signaling (Jeon et al., 2024), or phenylpropanoids as major components of plant defenses against pathogenic oomycetes (Carella et al., 2019). By contrast with these immune features conserved in tracheophytes and bryophytes, angiosperm specificities, such as the plasma membrane-located pattern recognition receptors FLS2 or EFR (Bowman et al., 2017), or the biosynthesis of Jasmonic Acid do not occur in bryophyte immunity (Monte et al., 2020), highlighting the independent evolution of defense mechanisms over the past 400 million years in the two main land plant lineages. So far, the discovery and identification of specific immune mechanisms has not been explored in bryophytes, with the noticeable exception of the defensive role played by oil-bodies against herbivores (Romani et al., 2020). Uncovering the origins and genetic basis of the bryophyte-specific immune mechanisms is essential to understanding how plant immune systems evolved across the green lineage. The liverwort *Marchantia polymorpha* offers unique advantages for such studies. This plant is a well-established model bryophyte, genetically tractable (Bowman et al., 2022), with a sequenced genome (Bowman et al., 2017) and a diversity of compatible interactions with well-described angiosperm pathogens including oomycete (Carella et al., 2018), bacteria (Gimenez-Ibanez et al., 2019), fungi (Matsui et al., 2020; Nelson et al., 2018) or virus (Ros-Moner et al., 2024). A collection of wild-collected accessions from Europe, the USA and Japan is also available for the three *M. polymorpha* subspecies (Beaulieu et al. 2024). Genomes of these accessions have been resequenced and compared, leading to the description of Single Nucleotide Polymorphisms (SNP) at the species level (Beaulieu et al. 2024), making *M. polymorpha* amenable for Genome-Wide Association Studies -GWAS- (Beaulieu et al. 2023).

Here, we leveraged these resources to investigate the genetic basis of quantitative resistance in *M. polymorpha* against the *M.* polymorpha-isolated fungal pathogen *Colletotrichum nymphaeae*. We first conducted complementary phenotypic, cytological and transcriptomic analyses to characterize a natural pathosystem between two *M. polymorpha* accessions with contrasting phenotypes following *C. nymphaeae* inoculation. Additionally, we utilized the genetic diversity within the *M. polymorpha* collection to perform the first Genome-Wide Association Study with a bryophyte. Our findings reveal both conserved and lineage-specific immune mechanisms, highlighting a potential role for terpene metabolism and horizontally transferred genes in the adaptation to this pathogen.

## Results

### Colletotrichum nymphaeae is a compatible parasite of Marchantia polymorpha Tak-1

To explore the genuine immune mechanisms of *M. polymorpha*, we selected one of the most pathogenic fungi identified in a previous survey of endophytes isolated from natural populations of *M. polymorpha* (Nelson et al., 2019). Previously referred to as *Colletotrichum sp*, we identified this strain as *C. nymphaeae* based on six classical makers used for *Colletotrichum sp*. taxonomy (Damm et al., 2014, see material and methods). To characterise the interaction between the model accession *M. polymorpha* Tak-1 and *C. nymphaeae*, a 10 µL droplet of conidia (10^4^/mL) was deposited in the centre of each thallus and a kinetics of infection was monitored over 6 days. The impact of *C. nymphaeae* on *M. polymorpha* Tak-1 development was monitored by scanning the plants every 24 hours, and comparing inoculated and mock-inoculated plants (Fig. 1). No symptoms were observed during the first two days following inoculation (Fig.1A). The first necrotic spots occurred at three days post inoculation (dpi) at the inoculation point (Fig. 1A), with brown tissues spreading throughout the thalli, resulting in fully macerated plants at 6 dpi (Fig. 1A). Meristematic tissues in some thalli remained green, an observation reminiscent of results described with angiosperm pathogens inoculated on *M. polymorpha* (Carella et al., 2018, Gimenez-Ibanez et al., 2019, Redkar et al., 2020). Differences between the thallus sizes measured before inoculation and at 6 dpi for each mock- or *C. nymphaeae*-inoculated thallus revealed a significant negative impact of *C. nymphaeae* on growth of *M. polymorpha Tak-1* thalli (Welch two-sample t.test, p-value = 6.436 x 10^-13^) (Fig. 1B).

**Figure 1.**
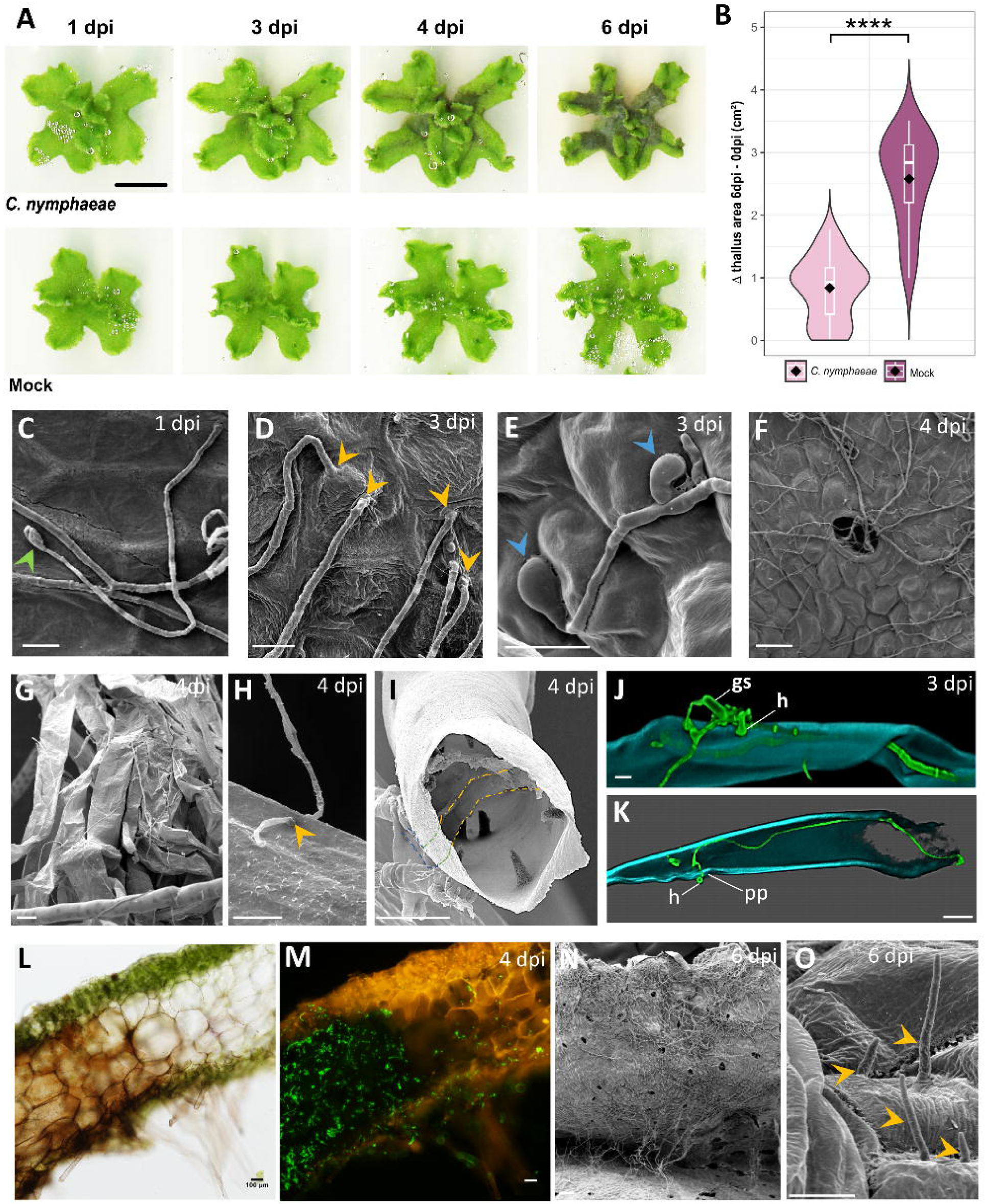
*C. nymphaeae* completes its biological cycle in *M. polymorpha*. **A** - Phenotypes of Tak-1 thalli from 0 to 6 dpi, after *C. nymphaeae* or mock inoculation (bar = 1cm). 40% of contrast and brightness increase from raw picture. **B** Boxplots and violin plots from n=68 measured samples. Each dot represents the difference between the surface of the thallus at 6 dpi and the measured surface of the thallus just before inoculation, for one sample, in inoculated (left) or mock (right) conditions. (****) indicates a p-value of 6.436e-13 from a Welch’s t-test **C to I** and **L** and **M**: Scanning Electron Microscopy (SEM) illustrating infection kinetics and specific features associated with *C. nymphaeae*’s biological cycle. **C**: Germinated spore (green arrowhead) at 1 dpi; **D**: Following germination, fungal hyphae generally directly penetrate thallus’ cuticle (yellow arrowheads) or, more rarely, (**E**) form appressorium-like structures (blue arrowheads) or (**F**) enter the thallus through airpores. (**C** to **E**; bar = 10µm; F: bar = 50µm). From 4 dpi, hyphae can also be easily observed, crawling on the rhizoids (**G, H** and **J**) or inside them, both in pegged (**I**) or in smooth (**K**) rhizoids, following a direct penetration (yellow arrowhead) (**H**) or using hyphopodia (**J** and **K**). Dashed lines (**I**) mark the hyphae outside (blue) or inside (green and yellow) the pegged rhizoid. **J** and **K** were obtained by confocal microscopy analyses following labelling of the mycelium with WGA-FITC (green) and of rhizoid cell walls by calcofluor (blue light). **gs**: germinated spore, **h**: hyphopodium, **pp**: penetration peg. (bars = 10 µm in **G**, **H** and **I** and 50 µm in **J** and **K**). At 4 dpi (**M**), hyphae decorated with WGA-FITC (in green) thrive in macerated and brown dead tissues (**L**). At 6 dpi, thalli are heavily colonised (**N**) and at this late time-point, *setae* (yellow arrowheads) emerged from the thalli (**O**). **L** and **M** is the same 100µm thin section of infected thallus observed 6 dpi in brightfield (**L**) or under excitation with blue light (M). (bars = 50µm in **L** and **M;** 100 µm in **N** and 10 µm in **O**).

Scanning Electronic Microscopy (SEM) was conducted on samples collected 2, 3, 4 and 6 dpi, to better describe the infection. Germination of the *C. nymphaeae* conidia typically occurred within the first day after inoculation (Fig. 1C). Fungal colonization was observed through diverse strategies, including direct penetration (Fig. 1D), development of appressorium-like structures (Fig. 1E), and colonization via air pores (Fig. 1F). At 4 dpi, the mycelium was clearly visible on the thallus surface (Fig. 1G, H, J), and inside the rhizoids (Fig. 1I, K). Rhizoid colonization occurs either directly (Fig. 1H) or via the formation of hyphopodia (Fig. 1J, K). Based on symptoms, the stage transition from biotrophic to necrotrophic takes place between 2 and 3 dpi. At 4 dpi fungal hyphae massively colonized brown, dead or dying cells (Fig. 1A, L, M). Finally, by 6 dpi the macerated thalli were heavily covered with mycelium (Fig. 1A, N). SEM analysis also revealed emerging setae, the fungal structures that precede the formation of the asexual fruiting body structures (acervuli) (Fig. 1O).

Altogether these results indicate that *C. nymphaeae* is a natural hemibiotrophic fungal parasite of *M. polymorpha*, capable of completing its biological cycle on the *M. polymorpha* Tak-1 accession during a fully compatible interaction.

### M. polymorpha displays quantitative resistance to Colletotrichum nymphaeae

Given that *C. nymphaeae* is a naturally occurring compatible pathogen of *M. polymorpha*, we hypothesized that diverse levels of resistance might exist across *M. polymorpha* populations. To test this hypothesis, we took advantage of the recently characterized collection of *M. polymorpha* accessions (Beaulieu et al., 2023). and selected a set of 87 easily propagated accessions to be assayed against *C. nymphaeae*. For each accession, 36 freshly collected gemmae were grown axenically on a sugar-free medium for three weeks, before being inoculated with *C. nymphaeae* or left untreated. Symptoms were monitored over time as described in the Methods section. Since the first symptoms are visible as early as 3 dpi in most accessions and showed little to no progression after 6 dpi, the latter timepoint was retained for symptoms quantification. The distribution of the symptom proportion (*i.e.* the ratio of symptomatic area - brown_area - to the total thallus area - thallus_area -) for the 87 accessions revealed a high variability of responses to *C. nymphaeae* (Fig. 2A). Indeed, the average proportion of symptoms per accession at 6 dpi ranged from 0 % to 59% for the most resistant and susceptible accessions, respectively. In the most resistant accessions, such as Nor-E, most thalli exhibit no symptoms at 6 dpi, suggesting complete resistance to *C. nymphaeae*. For other resistant accessions, such as CA, symptoms often remained restricted to the inoculation point in the middle of the thallus. By contrast, less resistant accessions displayed significant symptom development (Fig. S1). The reference accession *M. polymorpha* Tak-1 belongs to the group of most susceptible accessions (Fig. 2B).

**Figure 2.**
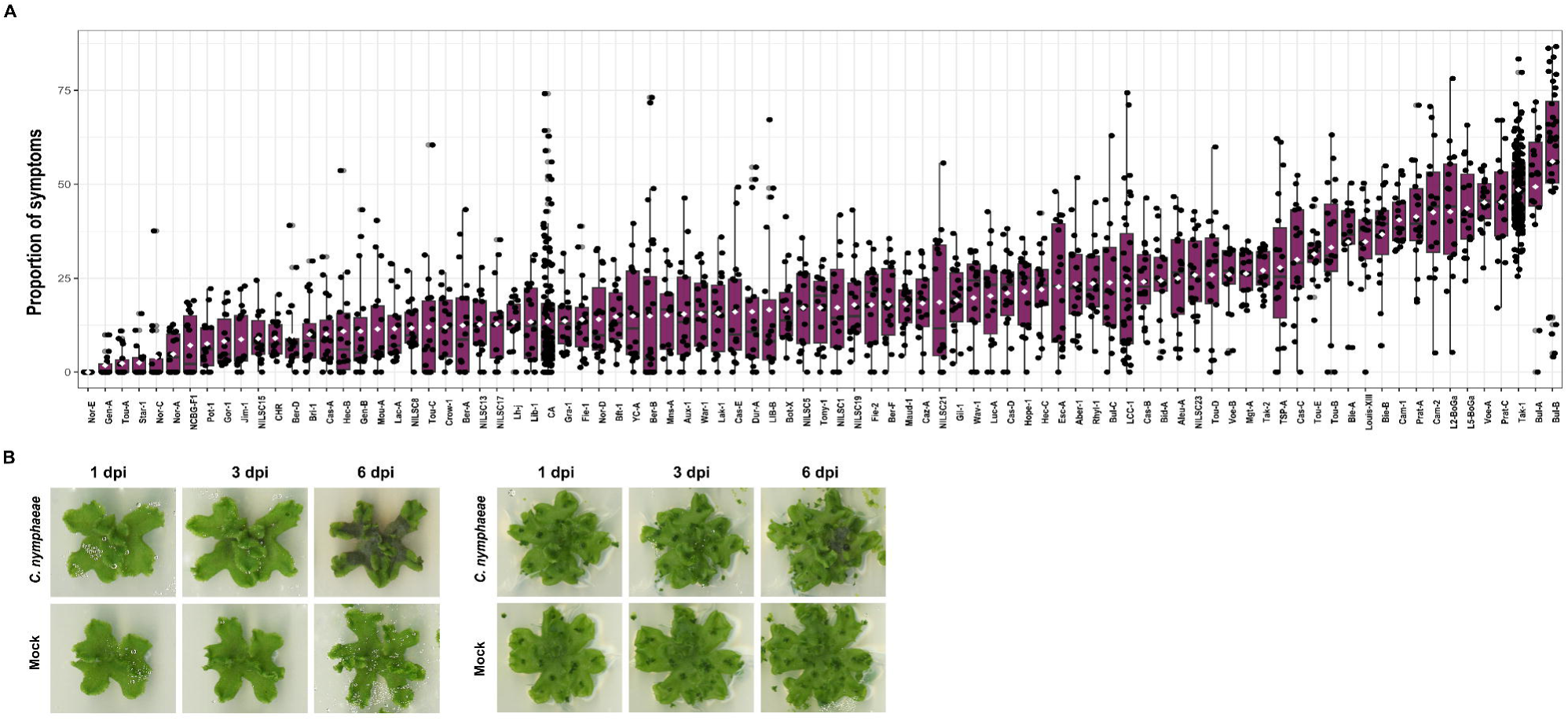
*M. polymorpha* exhibits quantitative resistance to *Colletotrichum nymphaeae*. A. Proportion of symptoms measured at 6 dpi for 87 accessions. For each accession, excluding the two internal controls Tak-1 and CA, 15 to 18 three weeks-old thalli were inoculated. B. Disease symptoms on *M. polymorpha* Tak-1 (left) and CA (right) thalli inoculated with *C. nymphaeae*, at 1, 3 and 6 dpi.

We conclude from this phenotypic screen that *M. polymorpha* exhibits quantitative resistance to *C. nymphaeae*.

### *Marchantia polymorpha* mounts a generic transcriptional response to filamentous pathogens

To investigate how *M. polymorpha* responds to infection by *C. nymphaeae*, we performed a RNAseq analysis on the susceptible accession *M. polymorpha* Tak-1 comparing the transcriptional profiles of 3-week-old thalli inoculated with water (mock) or with a suspension of *C. nymphaeae* for three and six days. Differential expression analysis (adjusted p-value ≤ 0.05 and absolute logFC [log2 fold change] ≥ 1) was conducted for mock-treated versus inoculated thalli at 3 dpi, as RNA extraction from 6 dpi was compromised due to excessive maceration. This analysis revealed strong transcriptional reprogramming in *M. polymorpha* Tak-1 following fungal infection (Fig. 3A, Data S1) with 1266 genes up-regulated and 575 genes down-regulated at 3 dpi. Functional enrichment analysis of *M. polymorpha* Tak-1 up-regulated genes identified InterPro domains associated with mechanical defenses, such as dirigent proteins and the O-methyltransferase COMT-type domains, which may be involved in cell wall-component biosynthesis. Domains associated with the specialized metabolism, including phenylpropanoids and flavonoid biosynthesis, with the phenylalanine ammonia-lyase and the chalcone/stilbene synthases domains, were also enriched (Fig. 3B). Additionally, domains implicated in general plant defenses and related signaling, such as chitinases, Bet v I (PR-proteins) and lipoxygenases (oxylipin pathway) were significantly enriched (Fig.3B, Data S2). Interestingly, genes most intensely up-regulated in response to *C. nymphaeae*, were predominantly accessory genes from the *M. polymorpha* ssp. *ruderalis* pangenome (63% of accessory genes in DEG with a logFC ≥5 vs 33% of accessory genes in DEG with a log FC ≤5, chi-square test *p-value* of 2.55x10^-12^). These results support the hypothesis that the accessory compartment of the pangenome plays an adaptive role in *M. polymorpha* immunity (Beaulieu et al., 2023). To determine whether this transcriptional reprogramming is specific to *C. nymphaeae* or part of a broader response, we compared our data with differential expression analyses of *M. polymorpha* infected by the oomycete *Phytophthora palmivora* (Carella et al., 2019), and by the bacterial pathogen *Pseudomonas syringae* DC3000 (Grenz et al., 2024). Among the genes up/down regulated in response to *C. nymphaeae* inoculation, 80%/3.4% and 57%/1.4% were also differentially regulated in Tak-1 following infections by *P. palmivora* / P*. syringae* respectively, suggesting a common up-regulation pattern in Marchantia, mainly in response to filamentous pathogens (Data S3). Notably, some Tak-1 genes up-regulated in response to *C. nymphaeae* (this study), to *P. palmivora* (Carella et al., 2019) and to *P. syringae* (Grenz et al., 2024) were also associated with climatic variation in a Genome-environment association study (Beaulieu et al., 2023). These include genes such as *MpNBS-LRR11* (Mp4g08790), *LURP1* (Mp4g08800) or *MpLOX5* (Mp1g21930), which were also differentially regulated in response to abiotic stresses (Data S4). This observation suggests their involvement in multiple stress responses, or reflect a close relationship between climatic conditions and pathogen pressures.

**Figure 3:**
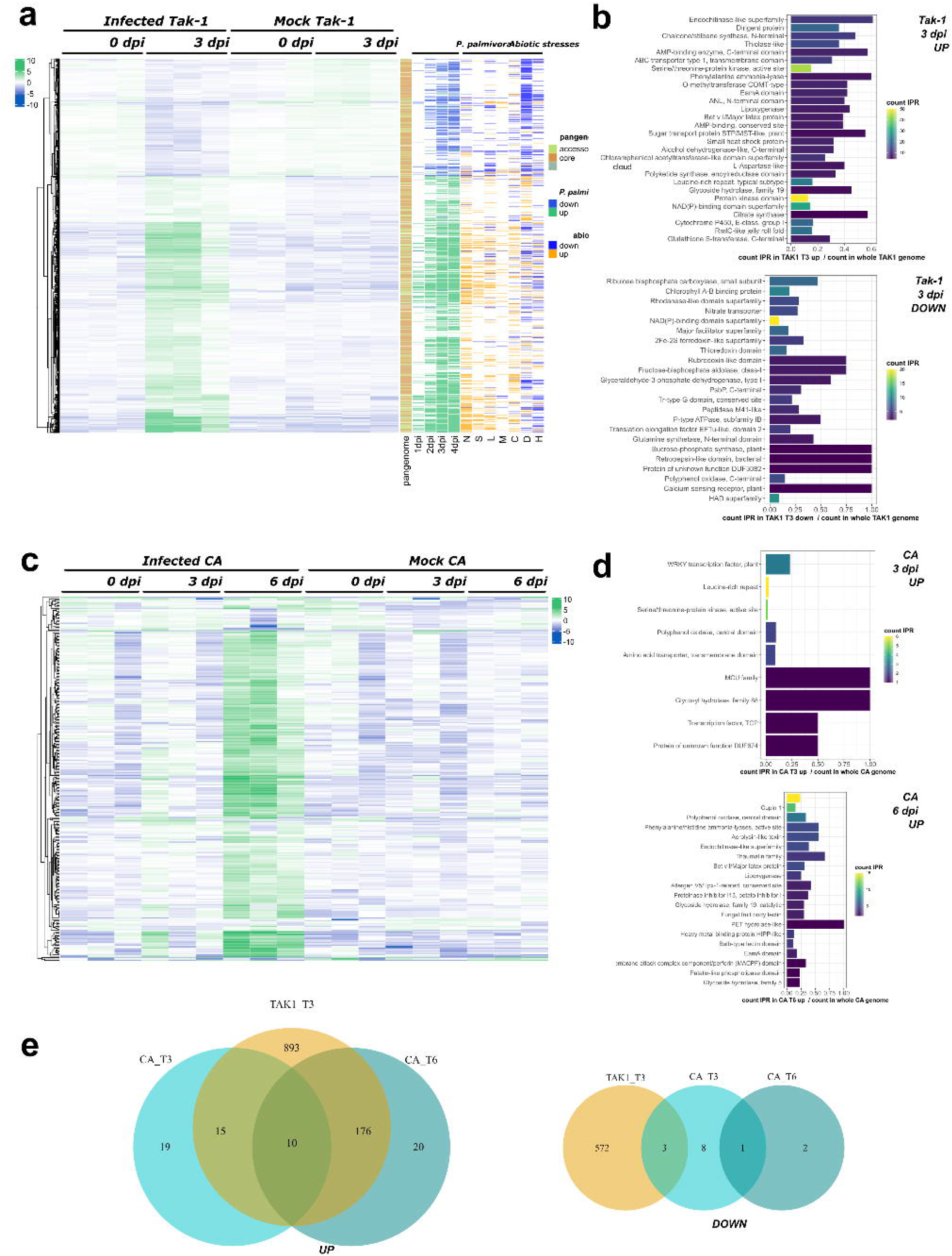
The *Marchantia polymorpha* ssp. *ruderalis* Tak-1 and CA accessions display striking temporal and functional discrepancies in their differential gene expression in response to *C. nymphaeae*. (a) Hierarchical clustering of Tak-1’s significantly differentially expressed genes during *C. nymphaeae* infection (adjusted p ≤ 0.05; log2 fold change [logFC] ≥ 1) at 0 and 3 days post inoculation (dpi). Variance stabilized row-centered counts are shown. Additional information on these genes is shown on the right side of the heatmap: pangenomic compartment of the genes, differential expression during the infection with *P. palmivora* (adjusted p-value ≤ 0.05; absolute log2 fold change [logFC] ≥ 1), and differential expression under different abiotic stresses (N=nitrogen deficiency, S=salt, L=light, M=mannitol, C=cold, D=dark, H=heat). (b) Functional enrichment of IPR terms in Tak-1’s up and down regulated genes at 3 dpi (enrichment cutoff 0.01). Redundant IPR terms were discarded to improve readability. (c) Hierarchical clustering of CA’s significantly differentially expressed genes during *C. nymphaeae* infection (adjusted p ≤ 0.05; absolute log2 fold change [logFC] ≥ 1) at 0, 3 and 6 days post inoculation (dpi). Variance stabilized row-centered counts are shown. (d) Functional enrichment of IPR terms in CAs up regulated genes at 3 and 6 dpi (enrichment cutoff 0.01). Redundant IPR terms were discarded to improve readability. (e) Comparisons of differentially regulated genes in Tak-1 and CA at different timepoints post inoculation (up regulated genes on the left side, down regulated genes on the right side). These Venn diagrams only represent the genes for which a gene-to-gene correspondence between CA and Tak-1 could be determined.

We conclude that the *M. polymorpha* Tak-1 transcriptional response to *C. nymphaeae* includes a generic response to microbial pathogens, enriched with pathogen-specific responses. This response partially overlaps, with general stress responses in *M. polymorpha*.

### Quantitative resistance is associated with a specific transcriptomic response

To assess the transcriptional differences between accessions with varying levels of resistance, a differential expression analysis was performed on one of the most resistant accession, *M. polymorpha* CA, isolated in the USA (Beaulieu et al., 2023), on samples collected at three and six dpi. At three dpi, only 44 genes were significantly up-regulated, and 12 down-regulated (Fig. 3C). At six dpi, 221 genes were significantly up-regulated and three genes down-regulated (adjusted p-value ≤ 0.05 and absolute logFC [log2 fold change] ≥ 1) (Fig. 3C, Data S5).

Compared to *M. polymorpha* Tak-1, the *M. polymorpha* CA accession exhibits a delayed transcriptional response to the pathogen, with only a few genes differentially expressed at 3 dpi (Fig. 3C), possibly indicating the presence of constitutive defenses in CA that delay the penetration of the pathogen and/or its perception by the plant. The limited transcriptional up-regulation at 3 dpi was enriched in domains from transcription factors (WRKY and TCP) and other signaling components (Leucine-rich repeats, kinases) commonly involved in plant – pathogen interactions (Fig. 3D, Data S6). Interestingly, one of the enriched domains, a glycosyl hydrolase 88 (GH88), is typically described in bacteria and fungi. Searching for this domain across plant lineages revealed its presence in mosses, liverworts, lycophytes and ferns, but not in hornworts nor in seed plants. Further phylogenetic analysis suggests a potential horizontal gene transfer (HGT) from fungi to the common ancestor of land plants, followed by subsequent losses in some lineages (Fig S2).

At 6 dpi, the transcriptional response of *M. polymorpha* CA becomes more pronounced, resembling the *M. polymorpha* Tak-1 response at 3 dpi. Enriched domains include dirigent proteins, PR10 (Bet v I/Major latex domain), lipoxygenases, phenylalanine ammonia lyases, polyphenol oxydases and chitinases. Other well-known defense-related protein families, such as various PR proteins (PR1: Allergen V5/Tpx-1-related, PR5: Thaumatin family, PR6: proteinase inhibitor, PR15: cupins) were also enriched. The Membrane Attack Complex Component/perforin (MACPF) domain, conserved across prokaryotes and eucaryotes was also present. MACPF proteins are involved in immune responses by forming pores in the membranes of pathogens (Rosado et al., 2008). Only two MACPF genes have been described in Arabidopsis and are involved in immune responses regulation in Arabidopsis (Morita-Yamamuro et al., 2005; Holmes et al., 2021). Additionally, fungal fruit body lectins originating from an HGT event in the ancestor of land plants (Beaulieu et al., 2023) are enriched in the *M. polymorpha* CA response to *C. nymphaeae*. To enable more precise comparison between the differentially expressed genes in the two accessions, a gene-to-gene correspondence was established between the two genomes (see Material and Methods). The overlap analysis (Fig. 3E) revealed ten genes commonly up-regulated at all timepoints in both accessions. These genes encode for proteins including transcription factors (MpWRKY3 and 7), secondary metabolite synthases (MpPAL7, MpPKS/CHS11, MpPPO3), peroxidase (PR9b), and an oxylipin synthesis enzyme (LOX1). Among the 18 genes uniquely up-regulated in CA, several encode proteins associated with Aerolisin/ETX pore-forming domain (4 genes), cupins (3 genes) and cytochrome P450 enzymes.

These results indicate a delayed response in *M. polymorpha* CA, mirroring the observed resistance, characterized by a minimal transcriptional reprogramming involving a core set of defense genes along with the activation of accession-specific genes.

### Identification of genetic loci associated with resistance/susceptibility variation by GWAS

Given the diversity of phenotypic responses to the inoculation by *C. nymphaeae* across the panel of *M. polymorpha* accessions, and the distinct transcriptional responses between the Tak-1 and CA accessions, we conducted a GWAS to explore the genetic bases of this responses. The association analysis was performed on 77 phenotyped accessions of *M. polymorpha ssp. ruderalis*, focusing on three phenotypes: the thallus area of inoculated plants at 6 dpi, the effect of the fungus on plant growth, and the area of symptoms (browning) at 6 dpi. Significant genomic loci were identified for each trait (Data S7).

In particular, the GWAS identified five genomic regions associated with symptom development (brown area), corresponding to 16 genes (Fig. 4). The peaks were distributed across autosomes. On chromosome 1, an association signal was found overlapping a methyltransferase and the functional regulatory region of a thioredoxin (Mp1g00650). Thioredoxins are involved in plant immunity through their reductase activity on cysteine residues that function as signalling switches, as well as their role in ROS detoxification (Mata-Pérez & Spoel, 2019). On chromosome 2, a strong association signal, was detected near genes encoding a tRNA synthetase (Mp2g20730) and a receptor-like kinase (Mp2g20720). At this locus, four of the most susceptible accessions share an alternative allele, potentially coupled with a small deletion, represented by missing SNP data (Fig. 4). This receptor-like kinase is the pro-ortholog of the 12 brassinosteroid signalling kinases from *A. thaliana* (Fig. 5), suggesting a possible role in brassinosteroid-mediated immunity, a pathway conserved across land plants (Yokota et al., 2017). Another association signal is present on chromosome 5, surrounded by two genes. One encodes an Orotidine 5’ decarboxylase (Mp5g15200) that usually participates in the synthesis of pyrimidine nucleotides. The other gene encodes an atypical kinase ABC1K (Mp5g15210), from a clade only found in algae, bryophytes and lycophytes, that has a common ancestor with the ABC1K12 mitochondrial family (Lundquist et al., 2012), whose role is not well defined. Interestingly, this gene is under selective sweep in Marchantia (top 14% of genes with low values of Fay and Wu’s *H,* Beaulieu et al. 2023).

**Figure 4:**
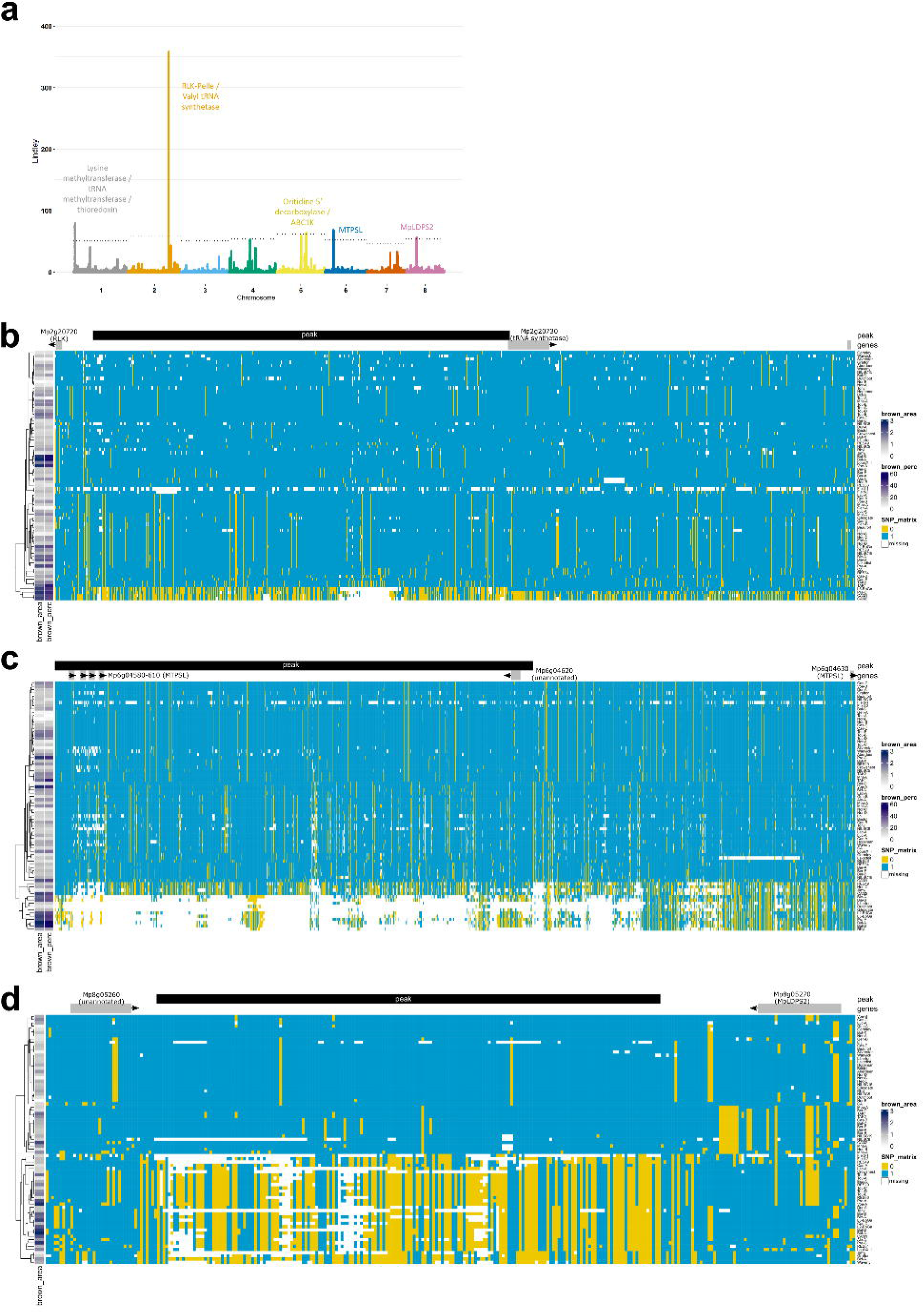
Identification of loci associated with phenotypic differences of *M. polymorpha* ssp *ruderalis* accessions in response to *C. nymphaeae*. (a) Manhattan plot of the GWAS results on the area of symptoms (brown area) on the thallus of inoculated plants at 6 dpi. This plot and the previous one result from a classical GWAS analysis performed with GEMMA, followed by the use of the local score technique on the SNP p-values, to amplify the signal between SNPs in LD. The result of this process is a Lindley value that has been used instead of p-value to plot the y values of the Manhattan plots. The dotted lines represent the significance thresholds for each chromosome (resampling thresholds from the local score method). (b) Haplotype block illustration of the genomic region on chromosome 2 associated with both the brown area of the symptoms (brown_area) and the ratio of symptoms on the total area of the thallus (brown_perc) in *M. polymorpha* ssp *ruderalis*. The association peak (black rectangle) is flanked by two protein-coding genes: an RLK-Pelle (Mp2g20720) and a tRNA synthetase (Mp2g20730). (c) Haplotype block illustration of the genomic region on chromosome 6 associated with both the brown area of the symptoms (brown_area) and the ratio of symptoms on the total area of the thallus (brown_perc) in *M. polymorpha* ssp *ruderalis*. The association peak (black rectangle) is overlapping 4 MTPSL genes (Mp6g04580-Mp6g04610) and an unannotated gene (Mp6g04620). A fifth MTPSL gene (Mp6g04630) is located downstream of the peak. (d) Haplotype block illustration of the genomic region on chromosome 8 associated with the brown area of the symptoms (brown_area) in *M. polymorpha* ssp *ruderalis*. The association peak (black rectangle) is flanked by two protein-coding genes: an unannotated gene (Mp8g05260) and the MpLDPS2 gene (Mp8g05270). For the 3 haplotype block illustrations, the gradient on the left side of the figure represents the values of the phenotypes of each accession that appear to be associated with the region represented. The main matrix represents the allelic status of the SNPs in this region for each accession: major allele (blue) or minor allele (yellow), missing information (white).

**Figure 5:**
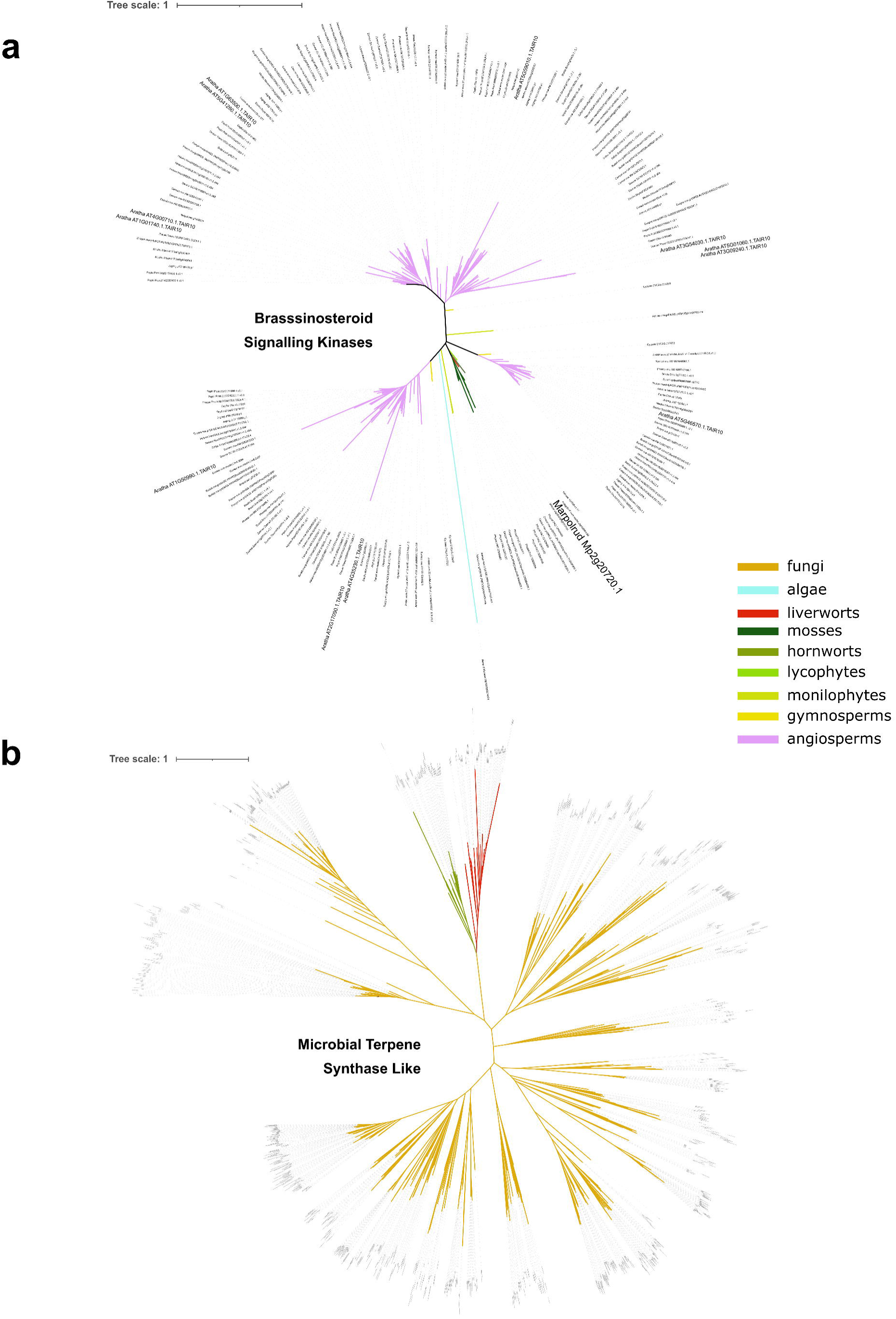
Phylogeny of the orthologs of the main GWAs candidate genes: the RLK Mp2g20720 and the 5 MTPSL Mp6g04580, Mp6g04590, Mp6g04605, Mp6g04610 and Mp6g04630. (a) Orthologs of the RLK GWAS candidate from chromosome 2 in viridiplantae. This tree was computed with the substitution model Q.plant+R7 and has a log-likelihood of -59286.8768. (b) Orthologs of the MTPSL GWAS candidates from chromosome 6 in viridiplantae and fungi. This tree was computed with the substitution model JTT+F+R10 and has a log-likelihood of - 271237.3539.

Further associations were found on chromosome 6 where a cluster of 5 microbial terpene synthase genes (MTPSL, Mp6g04580, Mp6g04590, Mp6g04605, Mp6g04610, Mp6g04630), is associated with variations in symptom severity. Some of the accessions with the largest area of browning in response to the pathogen display a common haplotype (bottom of the SNP matrix Fig. 4), with missing data in the SNP calling indicative of a potential deletion of this region in these accessions. Although none of the MTPSL-encoding gene from this cluster are deregulated in response to *C. nymphaeae*, the fifth one was up-regulated in response to *P. palmivora* (Carella et al.). This cluster of terpenes synthases, likely from the functional group III of monoterpene synthases (Kumar et al. 2016), belongs to one of the two families of microbial terpene synthases present in Marchantia, which is thought to have originated from a horizontal gene transfer from fungi (Jia et al., 2016). Phylogenetic analyses conducted on a much larger dataset than used previously (Jia et al. 2016) further confirms this hypothesis, and positions the HGT between an ancestor of the extant Dikarya fungi and the most recent common ancestor of the bryophytes (Fig. 5B). One of the drivers of tandem gene duplications is the transposon activity. To determine whether such a mechanism could have been involved in the evolution of the MTPSL cluster, we searched for signs of transposons in the surrounding genomic region. The same transposon residues were observed flanking each gene of the MTPSL cluster, suggesting that this family of 13 fungal terpene synthases was probably duplicated via transposon activity after its HGT from fungi. This mechanism could explain the substantial structural variation observed in this region of chromosome 6.

Another locus was found on chromosome 8, with an association region flanked by an unannotated gene (Mp8g05260) and the gene MpLDPS2 (Mp8g05270). Mp*LDPS2* encodes for a protein bearing a 1,8-cineole synthase domain, known to convert geranyl pyrophosphate, the precursor of monoterpenes, in 1,8-cineole (also known as eucalyptol). MpLDPS2 also contains a lipid droplet associated protein (LDPS) domain involved in the formation of lipid droplets. These structures not only contain lipids but also enzymes that enable the production of specialised metabolites, such as terpenes in bryophytes (de Vries & Ischebeck, 2020). Interestingly, the association peak on chromosome 8 is due to the balanced coexistence of two alleles, the minor allele being present in 34 accessions (*i.e.* 44% of all accessions studied), among which 16 are displaying severe browning symptoms (Fig. 4). The latter two candidate regions of the GWAS on the response of *M. polymorpha* to *C. nymphaeae*, the MTPSL and LDPS2, are pointing at a potential role of terpenes in the immune defenses of Marchantia, which is quite consistent with the known antimicrobial and antifungal effects of these specific metabolites (Asakawa & Ludwiczuk, 2018).

Together, these findings identified through the first GWAS analyses in bryophytes, suggest a complex genetic basis for quantitative resistance to *C. nymphaeae* in *M. polymorpha*, with key roles played by RLK, ROS homeostasis, cell wall reinforcement, proteasome regulation and secondary metabolism, including phenylpropanoid and terpene pathways, highlighting the importance of both core immune components and specialized metabolites.

## Discussion

Since the emergence of *M. polymorpha* as a suitable model for exploring evoMPMI (Schornak & Kamoun 2023) at the scale of land plants, most studies focused on describing the conservation of immune mechanisms (for review Delaux et Schornack, 2021; Castel *et al*., 2023; Ponce de León, 2023). However, lineage-specific immune mechanisms remain underexplored. To address this gap, we investigated the naturally-occurring pathosystem involving the filamentous hemibiotrophic pathogen *C. nymphaeae* and the genetic diversity within the *M. polymorpha* collection (Beaulieu et al., 2023). Our transcriptomic analysis revealed significant overlap with the response of *M. polymorpha* to *P. palmivora* infection, with 80% of the DEGs shared between the two datasets (Carella et al., 2019). This finding suggests a common response to filamentous pathogens and supports the conservation of immune mechanisms between *M. polymorpha* and angiosperms, as previously suggested, highlighting a core set of genes involved in plant defense. This includes PR proteins, enzymes of the phenylpropanoid pathway, and transcription factors (WRKY, NAC, AP2/ERF).

Many of the enriched genes families among the up-regulated genes such as LOX, NLR, and peroxidases, belong to the accessory genome of *M. polymorpha* (Beaulieu et al., 2023). This reinforces the proposed role of the accessory genome in adaptation to environmental stresses. Our analyses also identified accession-specific genes up-regulated in response to *C. nymphaeae*, but their precise role in defense against this pathogen requires further investigation.

Through GWAS, we identified terpene metabolism as a potential contributor to resistance to microbial infections in *M. polymorpha*. Although the underlying mechanisms remain to be validated, terpenes are well-known for their role in biotic stress responses (Chen et al., 2018). Liverworts, including *M. polymorpha*, exhibit a remarkable diversity of terpenes, with approximatively 1 600 identified compounds, including unique sesquiterpenes (Chen et al., 2018; Commisso et al., 2021). This diversity of terpenes is driven by terpene synthases (TPS) which produce a wide range of terpenes from common substrates (Karunanithi et Zerbe, 2019). *M. polymorpha* harbors both typical plant TPS and microbial-like TPS, the latter sharing similarities with bacterial and fungal enzymes (Jia et al., 2016). The dual presence of TPS types is characteristic of bryophytes and other non-seed plants, while MTPSL are absent in seed plants (Jia et al., 2016). The *M. polymorpha* Tak-1 genome encodes 32 MTPSL genes out of a total of 39 terpene synthase genes, including both bacterial-like MTPSL and fungal-like MTPSL genes (Bowman et al., 2017). In contrast, no MTPSL genes have been identified in the moss *P. patens* (Rensing et al., 2008), and the genomes of the sequenced hornworts (*A. agrestis* and *A. punctatus*) contain only 6 and 7 MTPSL genes, respectively (Li et al., 2020; Zhang et al., 2020). This variation may explain the observed differences in terpene diversity across bryophytes and other lineages, illustrating potential differences in their immune mechanisms. In *M. polymorpha*, terpenes accumulate in oil bodies that are liverworts-specific structures where MTPSL enzymes localize (Romani, Flores et al., 2022; Suire et al., 2000). These structures may also contribute to the large terpene diversity observed in liverworts. Although oil bodies contribute to defense against herbivores (Romani et al., 2020; Kanazawa et al., 2020), the molecules they store also exhibit antimicrobial properties (Romani, Flores et al., 2022), including antifungal activity (Chen et al., 2018). A potential role of oil body-stored terpenes in immunity, beyond herbivores, can be hypothesized.

MTPSL genes in *M. polymorpha* originated from horizontal gene transfer (HGT) events involving bacteria and fungi. A comprehensive analysis including bryophytes revealed two major HGT events involved in plant adaptation to terrestrial environments: one at the origin of streptophyte algae and another correlating with the plant terrestrialization. Interestingly, one-third of HGT-derived genes are associated with specialized metabolism (Ma, Wang et al., 2022), suggesting that MTPSL likely contributed to the adaptation to the terrestrial environments by enabling the synthesis of “specific” terpenes which may have been essential for defense. Beyond MTPSL, our analysis identified enrichment of FB lectin and Aerolysin/ETX pore-forming domains among the most significantly up-regulated genes in the *M. polymorpha* CA accession. These genes were also acquired through HGT (Ma, Wang *et al*., 2022) and may play roles immunity in *M. polymorpha*. For example, injection of FB lectin from *M. polymorpha* significantly increased the mortality rate of diamondback moth (Ma, Wang et al., 2022). A third example of HGT-acquired gene with relevance in plant defense is the GH88 domain, enriched among up-regulated genes in response to *C. nymphaeae*. Altogether, these findings suggest that HGT has significantly contributed to the evolution of the core land plants, and later on bryophyte-specific, immunity landscape. To conclude, the integration of transcriptomic analysis and GWAS identified core and Marchantia-specific immune mechanisms and emphasize the important roles of horizontal gene transfer and specialized metabolism in this process. Further investigation is required to understand the functions of these genes and loci in *M. polymorpha* immunity.

## Online methods

### Identification of *C. nymphaeae* growth and infection

*Colletotrichum spp.* strain used in this study was provided by J. Nelson (Nelson et al., 2018). Primers were designed to amplify six Colletotrichum spp. Sequences (Table S1), including *Glyceraldehyde 3-phosphate dehydrogenase* (*GAPDH*), Histone H3 (*HIS3*); Chitin synthase (*CHS-1*), *Actin* (*ACT*), *b-tubulin-2* (*TUB2*), and the internal transcribed spacers (ITS1 and ITS2), classically used to identify species of Colletotrichum species (Damm et al., 2014). The amplified fragments were Sanger sequenced and BLAST searches conducted against the nr database. For each sequence, the first hit value corresponded to the *C. nymphaeae* species with percentages of identity comprised between 92.2 to 99.49 % and e-value ranging from 5e^-113^ to 0.

### Phenotyping of various accessions in response to *C. nymphaeae*

Accessions of *Marchantia polymorpha* were cultivated from gemmae under axenic conditions and grown on ½ strength Gamborg B5 medium (Sigma, G5396), pH 5.7, 1,4% Agar-Agar (Sigma, A7921) under a long day photoperiod (16 h light at 22°C/8h dark at 20°C), with 60-80 μE light intensity. For each accession, to ensure the formation of uniform thalli 36 similar sized gemma were propagated, and distributed across four square plates (12*12 cm). From the four plates, two pairs were formed with each pair placed in a different phytotron that maintained identical conditions. After three weeks of development, one plate from each pair was used for inoculation, while the other remained unmanipulated to prevent contamination. Inoculation was performed by depositing a 10 µL droplet of suspension of *C. nymphaeae* (10^4^ spores/mL), at the base of each thallus. Following inoculation, the pairs of plates were placed back into their respective phytotrons. Symptoms were monitored daily through scans (EPSON GT20000). After each scan, the pairs were randomly rearranged within their phytotron.

Images were analyzed using the software Image Pro Plus © for the studied timepoints. A macro (Data S8) was used to automate the analysis and determine the thallus area. Symptomatic areas were manually adjusted by applying “Split Object” or “Draw/Merge Objects”.

The *Marchantia polymorpha* collection was split into eight experimental batches during phenotyping, and two accessions were used as internal controls: CA and Tak-1, respectively resistant and susceptible to *C. nymphaeae*. Two weeks before inoculation, 200 µI of a 50% glycerol stock were spread on Mathur medium. Spores were collected by adding 5 mL of sterile water and scraping the mycelium. Infection experiments were performed by applying two 10 μL droplets of a 10^4^ spores/mL suspension at the base of the thallus.

### Scanning Electron Microscopy (SEM)

Samples were fixed in a 0.05 M sodium cacodylate solution (pH 7.2) containing 2.5% glutaraldehyde, then dehydrated through a graded ethanol series. Critical-point drying was performed using liquid CO₂. The samples were mounted on an observation plate with the rhizoids facing upwards and grounded using conductive silver paint. They were sputter-coated with platinum before imaging. Images were acquired using a scanning electron microscope (Quanta 250 FEG, FEI) at 5 kV, with a spot size of 3 and a working distance of 10 mm.

### Bright field microscopy

For optical microscopy, 110-µm-thick sections of fresh *M. polymorpha* thalli were collected at various time points after inoculation with *C. nymphaeae*. Thalli were embedded in 5% low– melting-point agarose and sectioned using a vibratome (VT1000S; Leica, Rueil-Malmaison, France). The thallus sections, including rhizoids, were stained with WGA-FITC (50 µg/mL in PBS), washed three times with PBS, and mounted on glass slides in a drop of PBS. Observations were performed using bright-field microscopy or epifluorescence microscopy with an inverted microscope (Nikon Eclipse TI) equipped with a color CMOS camera DS Ri2 and controlled by Nikon NIS software. A 10x (N.A 0.3) objective was used. The fluorescence of WGA-FITC was observed using a GFP SMO filter set (excitation: 472/30 nm; dichroic: 495 nm; emission: 520/35 nm), while the autofluorescence of the cell walls was detected using a CY3 filter set (excitation: 543/22 nm; dichroic: 560 nm; emission: 590/40 nm). Alternatively, samples were analyzed using a confocal microscope (LEICA SP8) at the FRAIB imaging platform.

### Confocal microscopy

Confocal images of rhizoids and *Colletotrichum* were acquired using a spectral confocal laser scanning system (SP8, Leica, Germany) equipped with an upright microscope (DMi8, Leica, Germany). Observations were performed with a 10× dry objective (HC PL FLUOTAR, N.A. 0.30). An Argon laser emitting at 488 nm was used to detect WGA-FITC fluorescence in the range of 500–540 nm. Additionally, a laser diode emitting at 405 nm was used to visualize rhizoid cell walls stained with calcofluor, with fluorescence collected in the range of 410–460 nm. Images were acquired in sequential mode to prevent inter-channel crosstalk. Fluorescence overlay images were generated by projecting 20–30 confocal planes acquired along the z-axis with a 4-µm increment between focal planes, creating a 3D projection.

### RNA-Seq experiment

Three weeks-old thalli from CA and Tak-1 were mock inoculated (water) or with a suspension of *C. nymphaeae*. For each kinetic timepoint, the meristematic zones were harvested separately from the rest of the thallus, as these zones are symptom free and believed to exhibit a distinct response to infection. The samples were then frozen in liquid nitrogen, grounded and stored at -70°C until RNA extraction. Total RNA was extracted from ∼100 mg of grounded thalli using the DirectZol kit (Ozyme, R2052) following the manufacturer’s instructions and sent for sequencing to Genewiz (Leipzig, Germany). Illumina librairies were prepared and sequenced on a NovaSeq platform using 150 bp paired-end reads, generating approximately 20 million reads per sample.

### Annotation of the CA genome

The genome from CA (Beaulieu et al. 2023) was soft-masked using Earlgrey v4.3.0 (Baril et al., 2024) and the structural annotation was conducted using BRAKER v3.0.7 pipeline (Brůna et al., 2020; Gabriel et al., 2021, 2023; Kovaka et al., 2019; Pertea & Pertea, 2020; Quinlan, 2014; Stanke et al., 2006, 2008). BRAKER2 was run with –prot_seq –bam --gff3 and – busco_lineage options. In ETPmode the GeneMark-ETP pipeline generates hints based on the alignment of RNAseq reads and on the protein database supplied which enables AUGUSTUS training. The prediction of protein coding genes is a combination of AUGUSTUS and GeneMark-ETP predictions. The OrthoDB input proteins used by ProtHint is a combination of https://v100.orthodb.org/download/odb10_plants_fasta.tar.gz and proteins from seven species (Anthoceros agrestis cv. BONN, Anthoceros agrestis cv. OXF, Anthoceros punctatus (F.-W. Li et al., 2020), Ceratodon purpureus strain R40 (NCBI GCA_014871385.1), Marchantia paleacea (Radhakrishnan et al., 2020), Marchantia polymorpha ssp. ruderalis TAK1 (https://marchantia.info/download/MpTak_v6.1/), Physcomitrium patens (Lang et al., 2018) and Sphagnum fallax (Healey et al., 2023)). The completeness of the prediction was assessed with Compleasm (Huang & Li, 2023) against the viridiplantae odb10 (n=425). The predictions were functionally annotated with interproscan-5.64-96.0 (Blum et al., 2021; Jones et al., 2014) with options –iprlookup and – goterms. The gene-to-gene correspondence between Tak-1’s and CA’s genome was determined combining the information from a collinearity comparison of the two genomes with MCscanX (Y. Wang et al., 2012) and the mapping of Tak-1’s annotation on CA’s genome with Liftoff v1.6.3 (Shumate & Salzberg, 2021).

### Expression analysis

The raw reads were processed and mapped to their representative genome (Marchantia polymorpha Tak1 v6 and Marchantia polymorpha CA v1) with nextflow v21.10.6 (Di Tommaso et al., 2017) and the nf-core/rnaseq r3.9 (Ewels et al., 2020) pipeline, using the -- skip_qc --aligner star_salmon --remove_ribo_rna options. The workflow used bedtools v2.30.0 (Quinlan, 2014), gffread v0.12.1 (Pertea & Pertea, 2020), star v2.7.10a (Dobin et al., 2013), picard v2.27.4, salmon v1.5.2 (Patro et al., 2017), SummarizedExperiment v1.20.0 (Morgan et al., 2024), tximeta v1.8.0 (Love *et al*., 2020), samtools v1.15.1 (H. Li et al., 2009), sortmerna v4.3.4 (Kopylova et al., 2012), stringtie v2.2.1 (Kovaka et al., 2019), trimgalore v0.6.7 and usc v377. Differentially expressed genes were identified using the edgeR package (Robinson et al., 2010) in R v4.4.0, separately on each accession. Low-expressed genes with less than ten reads across each class of samples were removed and gene counts were normalized by library size and using the trimmed mean of M-values normalization method (Robinson & Oshlack, 2010). DEGs were estimated by pairwise comparisons between infected and mock inoculated samples at the same stage of infection. Differentially expressed genes (adjusted p-value ≤ 0.05 and absolute logFC [log2 fold change] ≥ 1) were used to perform hierarchical clustering of samples. Heatmaps for the differentially expressed genes were generated using R ComplexHeatmap package (Gu et al., 2016) using variance-stabilized counts median-centered by gene.

### IPR term enrichment

Enrichment analyses were performed on the different categories of differentially expressed genes (in each accession, at each timepoint, up and down regulated genes), except from the ones that had a low number of differentially expressed gene (0 dpi, down regulated in CA). The enrichment was performed with the FUNC-E package v2.0.1 (Ficklin et al., 2021) with an enrichment cut of 0.01.

### Estimation of accessions phenotypic means

Phenotypic data on the thallus area, browning area of the thallus, and on the ratio between both, in inoculated plants at 0 and 6 dpi, were used to perform a genome wide association study. First, for each accession, outlier individuals (with their value being higher or lower than 1.5 times the interquartile range) for at least two phenotypic variables were discarded. Then, a linear model was applied on the data for each phenotypic variable (except the thallus area pre-inoculation, that was used as a covariable) in order to estimate the accessions means according to various confounding effects (effect of the experimental batch, of the phytotron and of the area of the thallus pre-inoculation). For the Tak-1 (susceptible) and CA (resistant) accessions that were present in each experimental batch, the linear model is as follows: *phenotype_ijkl_ = µ + accession_i_ + phytotron_j_ * batch_k_ + preinoc_thallus_area_l_ + E_ijkl_*, where *µ* is the overall mean, “accession” corresponds to the difference between Tak-1 and CA accessions, “phytotron” and “batch” account for the effect of the two phytotrons used and the experimental batch in which each plant was grown, “preinoc_thallus_area” is a quantitative covariate accounting for the initial thallus size of each plant, and E is the residual term. This allows to get adjusted means for CA and Tak-1, that will be used as their phenotypic values in the GWAS. In order to use Tak-1 as an internal control of the experimental batches involving all other *M. polymorpha* accessions, another linear model is implemented that only considers the experimental batch and the initial thallus size of each Tak-1 plant before inoculation: *phenotype_ij_ =µ + batch_i_ + preinoc_thallus_area_j_ + E_ij_*. The estimated “batch” effects are then used as a covariable (TAK1_ctrl) for the other accessions in the same experimental batch, in the following linear model: *phenotype_ijkl_ =µ + accession_i_ * phytotron_j_ + TAK1_ctrl_k_ + preinoc_thallus_area_l_ + E_ijkl_*. For each phenotype analysed, adjusted means of each *M. polymorpha* accession is estimated, and then serve as inputs in the GWAS.

### Genome wide association study

GWAS analyses were performed using the mixed linear model implemented in GEMMA software (v 0.98.1 (Zhou & Stephens, 2012)) on a dataset of 2 141 087 SNPs with minor allele frequencies of 0.05 and maximum of 15 accessions with missing data per site, on a set of 77 phenotyped accessions from the subspecies *ruderalis*. To estimate the SNPs effects and their significance, the model used a centered kinship matrix as a covariable with random effect, and a Wald test. SNP *P*-values from GEMMA were processed using a local score approach (Bonhomme et al., 2019; Fariello et al., 2017) to help detect robust loci across analyses. The local score is a cumulative score that takes advantage of local linkage disequilibrium among SNPs. This score, defined as the maximum of the Lindley process over a SNP sequence (i.e., a chromosome), was calculated using a tuning parameter value of ξ=2, as suggested by simulation results (Bonhomme et al., 2019). Chromosome-specific significance thresholds (C =5%) were estimated using a resampling approach. The R scripts used to compute the local score and significance thresholds are available at https://forge-dga.jouy.inra.fr/projects/local-score/documents. All the significant local score peaks coordinates were extracted, and they were annotated with their overlapping (when existing), downstream and upstream genes.

### Phylogenies of candidate genes

Phylogeny for the MTPSL genes (Mp6g04580, Mp6g04590, Mp6g04605, Mp6g04610, Mp6g04630) was determined by BLASTp+ v2.12.0 (Camacho et al., 2009) (maximum of 2000 target sequences and e-value of 10^-5^) against a database of viridiplantae genomes (Data S9), a database with non-angiosperm transcriptomes from the 1KP initiative (One Thousand Plant Transcriptomes Initiative, 2019), a database with fungal genomes from MycoCosm [(Grigoriev et al., 2014) last time consulted 02/2019], and the non-redundant (nr) database from the NCBI. The resulting proteins were aligned with muscle5 v5.1 (Edgar, 2022) and trimmed with trimAl v1.4 (Capella-Gutierrez et al., 2009) to discard positions with more than 60% of gaps. The phylogenetic tree was computed with IQ-TREE v2.1.2 and the best-fitting evolutionary model selected using modelFinder according to the Bayesian Information Criteria. Branch support was estimated with 10 000 replicates of both SH-like approximate likelihood ratio and ultrafast bootstrap. For the receptor-like kinase (Mp2g20720), the same tools were used but the sequence research was only performed against the custom database of plant genomes. For the GH88 domain, sequence research and alignment were performed the same way, but the phylogeny was performed with FastTree v2.1.11 (Price et al., 2010) with the default options.

## Acknowledgments

The authors thank the genotoul bioinformatics platform Toulouse Occitanie (Bioinfo Genotoul, https://doi.org/10.15454/1.5572369328961167E12) for providing computing resources. Research at the LRSV is supported by the Laboratoires d’Excellence (LABEX)’ TULIP (ANR-10-LABX-41). This project received funding from the European Research Council (ERC) under the European Union’s Horizon 2020 research and innovation programme (grant agreement no. 101001675 - ORIGINS) and from the FRM/FSER (FRM/FSER202302017064) to P.-M.D, from the CNRS to P-M.D (80|PRIME MicMac), from the Agence Nationale de la Recherche (ANR LEVEL-UP ANR-21-CE20-0010-01) to C.J.

## Author contributions

K.E.M, B.C., N.J., E.A., F.B., Y.M., J.N. carried out experiments. K.E.M., C.B., C.L., F.B., J.K., M.B. analysed data. K.E.M., C.B., M.B., C.J., and P-M.D wrote the manuscript. M.B., C.J., P-M.D coordinated the project.

## References

Asakawa, Y., & Ludwiczuk, A. (2018). Chemical Constituents of BryophytesC: Structures and Biological Activity. Journal of Natural Products, 81(3), 641–660. 10.1021/acs.jnatprod.6b01046

Baril, T., Galbraith, J., & Hayward, A. (2024). Earl GreyC: A Fully Automated User-Friendly Transposable Element Annotation and Analysis Pipeline. Molecular Biology and Evolution, 41(4), msae068. 10.1093/molbev/msae068

Beaulieu, C., Libourel, C., Zamar, D. L. M., Mahboubi, K. E., Hoey, D. J., Keller, J., Girou, C., Clemente, H. S., Diop, I., Amblard, E., Théron, A., Cauet, S., Rodde, N., Zachgo, S., Halpape, W., Meierhenrich, A., Laker, B., Brautigam, A., Greiff, G. R., … Delaux, P. (2023). TheMarchantiapangenome reveals ancient mechanisms of plant adaptation to the environment. bioRxiv (Cold Spring Harbor Laboratory). 10.1101/2023.10.27.564390

Beerling, D. (2007). The Emerald Planet: How plants changed Earth’s history. The Emerald 585 Planet. 10.1093/OSO/9780192806024.001.0001.

Blum, M., Chang, H.-Y., Chuguransky, S., Grego, T., Kandasaamy, S., Mitchell, A., Nuka, G., Paysan-Lafosse, T., Qureshi, M., Raj, S., Richardson, L., Salazar, G. A., Williams, L., Bork, P., Bridge, A., Gough, J., Haft, D. H., Letunic, I., Marchler-Bauer, A., … Finn, R. D. (2021). The InterPro protein families and domains databaseC: 20 years on. Nucleic Acids Research, 49(D1), D344–D354. 10.1093/nar/gkaa977

Bonhomme, M., Fariello, M. I., Navier, H., Hajri, A., Badis, Y., Miteul, H., Samac, D. A., Dumas, B., Baranger, A., Jacquet, C., & Pilet-Nayel, M.-L. (2019). A local score approach improves GWAS resolution and detects minor QTLC: Application to Medicago truncatula quantitative disease resistance to multiple Aphanomyces euteiches isolates. Heredity, 123(4), 517–531. 10.1038/s41437-019-0235-x

Bowman, J. L., Kohchi, T., Yamato, K. T., Jenkins, J., Shu, S., Ishizaki, K., Yamaoka, S., Nishihama, R., Nakamura, Y., Berger, F., Adam, C., Aki, S. S., Althoff, F., Araki, T., Arteaga-Vazquez, M. A., Balasubrmanian, S., Barry, K., Bauer, D., Boehm, C. R., … Schmutz, J. (2017). Insights into Land Plant Evolution Garnered from the Marchantia polymorpha Genome. Cell, 171(2), 287–304.e15. 10.1016/j.cell.2017.09.030

Bowman, J. L., Arteaga-Vazquez, M., Berger, F., Briginshaw, L. N., Carella, P., Aguilar-Cruz, A., Davies, K. M., Dierschke, T., Dolan, L., Dorantes-Acosta, A. E., Fisher, T. J., Flores-Sandoval, E., Futagami, K., Ishizaki, K., Jibran, R., Kanazawa, T., Kato, H., Kohchi, T., Levins, J., … Zachgo, S. (2022). The renaissance and enlightenment ofMarchantiaas a model system. The Plant Cell, 34(10), 3512–3542. 10.1093/plcell/koac219

Bressendorff, S., Azevedo, R., Kenchappa, C.S., Ponce de León, I., Olsen, J.V., Rasmussen, M.W., Erbs, G., Newman, M.-A., Petersen, M., and Mundy, J. (2016). An Innate Immunity Pathway in the Moss Physcomitrella patens. The Plant Cell 28, 1328–1342. 10.1105/tpc.15.00774.

Brůna, T., Lomsadze, A., & Borodovsky, M. (2020). GeneMark-EP+C: Eukaryotic gene prediction with self-training in the space of genes and proteins. NAR Genomics and Bioinformatics, 2(2), lqaa026. 10.1093/nargab/lqaa026

Camacho, C., Coulouris, G., Avagyan, V., Ma, N., Papadopoulos, J., Bealer, K., & Madden, T. L. (2009). BLAST+C: Architecture and applications. BMC Bioinformatics, 10(1), 421. 10.1186/1471-2105-10-421

Capella-Gutierrez, S., Silla-Martinez, J. M., & Gabaldon, T. (2009). trimAlC: A tool for automated alignment trimming in large-scale phylogenetic analyses. Bioinformatics, 25(15), 1972–1973. 10.1093/bioinformatics/btp348

Carella, P., Gogleva, A., Hoey, D. J., Bridgen, A. J., Stolze, S. C., Nakagami, H., & Schornack, S. (2019). Conserved Biochemical Defenses Underpin Host Responses to Oomycete Infection in an Early-Divergent Land Plant Lineage. Current Biology, 29(14), 2282–2294.e5. 10.1016/j.cub.2019.05.078

Carella, P., Gogleva, A., Tomaselli, M., Alfs, C., & Schornack, S. (2018). *Phytophthora palmivora* establishes tissue-specific intracellular infection structures in the earliest divergent land plant lineage. Proceedings of the National Academy of Sciences, 115(16). 10.1073/pnas.1717900115

Castel, B., El Mahboubi, K., Jacquet, C., & Delaux, P. (2023). Immunobiodiversity: conserved and specific immunity across land plants and beyond. Molecular Plant, 17(1), 92–111. 10.1016/j.molp.2023.12.005

Chen, F., Ludwiczuk, A., Wei, G., Chen, X., Crandall-Stotler, B., and Bowman, J.L. (2018). Terpenoid Secondary Metabolites in Bryophytes: Chemical Diversity, Biosynthesis and Biological Functions. Critical Reviews in Plant Sciences 37, 210 231. 10.1080/07352689.2018.1482397.

Commisso, M., Guarino, F., Marchi, L., Muto, A., Piro, A., and Degola, F. (2021). Bryo-Activities: A Review on How Bryophytes Are Contributing to the Arsenal of Natural Bioactive Compounds against Fungi. Plants 10, 203. 10.3390/plants10020203.

Damm U, O’Connell RJ, Groenewald JZ, Crous PW (2014) The Colletotrichum destructivum species complex - hemibiotrophic pathogens of forage and field crops. Stud Mycol 79: 49–84

Delaux, P.-M., Hetherington, A. J., Coudert, Y., Delwiche, C., Dunand, C., Gould, S., Kenrick, P., Li, F.-W., Philippe, H., Rensing, S. A., Rich, M., Strullu-Derrien, C., & de Vries, J. (2019). Reconstructing trait evolution in plant evo–devo studies. Current Biology, 29(21), R1110–R1118. 10.1016/j.cub.2019.09.044

Delaux, P.-M., & Schornack, S. (2021). Plant evolution driven by interactions with symbiotic and pathogenic microbes. Science, 371(6531), eaba6605. 10.1126/science.aba6605

de Vries, J., & Ischebeck, T. (2020). Ties between Stress and Lipid Droplets Pre-date Seeds. Trends in Plant Science, 25(12), 1203–1214. 10.1016/j.tplants.2020.07.017

Di Tommaso, P., Chatzou, M., Floden, E. W., Barja, P. P., Palumbo, E., & Notredame, C. (2017). Nextflow enables reproducible computational workflows. Nature Biotechnology, 35(4), 316–319. 10.1038/nbt.3820

Dobin, A., Davis, C. A., Schlesinger, F., Drenkow, J., Zaleski, C., Jha, S., Batut, P., Chaisson, M., & Gingeras, T. R. (2013). STARC: Ultrafast universal RNA-seq aligner. Bioinformatics, 29(1), 15–21. 10.1093/bioinformatics/bts635

Edgar, R. C. (2022). Muscle5C: High-accuracy alignment ensembles enable unbiased assessments of sequence homology and phylogeny. Nature Communications, 13(1), 6968. 10.1038/s41467-022-34630-w

Ewels, P. A., Peltzer, A., Fillinger, S., Patel, H., Alneberg, J., Wilm, A., Garcia, M. U., Di Tommaso, P., & Nahnsen, S. (2020). The nf-core framework for community-curated bioinformatics pipelines. Nature Biotechnology, 38(3), 276–278. 10.1038/s41587-020-0439-x

Fariello, M. I., Boitard, S., Mercier, S., Robelin, D., Faraut, T., Arnould, C., Recoquillay, J., Bouchez, O., Salin, G., Dehais, P., Gourichon, D., Leroux, S., Pitel, F., Leterrier, C., & SanCristobal, M. (2017). Accounting for linkage disequilibrium in genome scans for selection without individual genotypesC: The local score approach. Molecular Ecology, 26(14), 3700–3714. 10.1111/mec.14141

Ficklin, S., Bensman, E., Poehlman, W., & Feltus, A. (2021). SystemsGenetics/FUNC-E: (Version v2.0.0) [Logiciel]. Zenodo. 10.5281/ZENODO.5587344

Gabriel, L., Hoff, K. J., Brůna, T., Borodovsky, M., & Stanke, M. (2021). TSEBRAC: Transcript selector for BRAKER. BMC Bioinformatics, 22(1), 566. 10.1186/s12859-021-04482-0

Gabriel, L., Brůna, T., Hoff, K. J., Ebel, M., Lomsadze, A., Borodovsky, M., & Stanke, M. (2023). BRAKER3J: Fully automated genome annotation using RNA-seq and protein evidence with GeneMark-ETP, AUGUSTUS and TSEBRA. 10.1101/2023.06.10.544449

Gimenez-Ibanez, S., Zamarreño, A. M., García-Mina, J. M., & Solano, R. (2019). An Evolutionarily Ancient Immune System Governs the Interactions between Pseudomonas syringae and an Early-Diverging Land Plant Lineage. Current Biology, 29(14), 2270–2281.e4. 10.1016/j.cub.2019.05.079

Grenz, K., Chia, K., Turley, E. K., Tyszka, A. S., Atkinson, R. E., Reeves, J., Vickers, M., Rejzek, M., Walker, J. F., & Carella, P. (2024). A Necrotizing Toxin Promotes Pseudomonas syringae Infection Across Evolutionarily Divergent Plant Lineages. bioRxiv (Cold Spring Harbor Laboratory). 10.1101/2024.07.17.603760

Grigoriev, I. V., Nikitin, R., Haridas, S., Kuo, A., Ohm, R., Otillar, R., Riley, R., Salamov, A., Zhao, X., Korzeniewski, F., Smirnova, T., Nordberg, H., Dubchak, I., & Shabalov, I. (2014). MycoCosm portalC: Gearing up for 1000 fungal genomes. Nucleic Acids Research, 42(D1), D699–D704. 10.1093/nar/gkt1183

Gu, Z., Eils, R., & Schlesner, M. (2016). Complex heatmaps reveal patterns and correlations in multidimensional genomic data. Bioinformatics, 32(18), 2847–2849. 10.1093/bioinformatics/btw313

Han, G. (2019). Origin and evolution of the plant immune system. New Phytologist 222, 70–83. 10.1111/nph.15596.

Healey, A. L., Piatkowski, B., Lovell, J. T., Sreedasyam, A., Carey, S. B., Mamidi, S., Shu, S., Plott, C., Jenkins, J., Lawrence, T., Aguero, B., Carrell, A. A., Nieto-Lugilde, M., Talag, J., Duffy, A., Jawdy, S., Carter, K. R., Boston, L.-B., Jones, T., … Shaw, A. J. (2023). Newly identified sex chromosomes in the Sphagnum (peat moss) genome alter carbon sequestration and ecosystem dynamics. Nature Plants, 9(2), 238–254. 10.1038/s41477-022-01333-5

Holmes, D. R., Bredow, M., Thor, K., Pascetta, S. A., Sementchoukova, I., Siegel, K. R., Zipfel, C., & Monaghan, J. (2021). A novel allele of the Arabidopsis thaliana MACPF protein CAD1 results in deregulated immune signaling. Genetics, 217(4). 10.1093/genetics/iyab022

Huang, N., & Li, H. (2023). compleasmC: A faster and more accurate reimplementation of BUSCO. Bioinformatics, 39(10), btad595. 10.1093/bioinformatics/btad595

Jeon HW, Iwakawa H, Naramoto S, Herrfurth C, Gutsche N, Schlüter T, Kyozuka J, Miyauchi S, Feussner I, Zachgo S, Nakagami H. Contrasting and conserved roles of NPR pathways in diverged land plant lineages. New Phytol. 2024 Sep;243(6):2295–2310. doi: 10.1111/nph.19981. Epub 2024 Jul 26. PMID: 39056290.

Jia, Q., Li, G., Köllner, T. G., Fu, J., Chen, X., Xiong, W., Crandall-Stotler, B. J., Bowman, J. L., Weston, D. J., Zhang, Y., Chen, L., Xie, Y., Li, F.-W., Rothfels, C. J., Larsson, A., Graham, S. W., Stevenson, D. W., Wong, G. K.-S., Gershenzon, J., & Chen, F. (2016). Microbial-type terpene synthase genes occur widely in nonseed land plants, but not in seed plants. Proceedings of the National Academy of Sciences, 113(43), 12328–12333. 10.1073/pnas.1607973113

Jones, P., Binns, D., Chang, H.-Y., Fraser, M., Li, W., McAnulla, C., McWilliam, H., Maslen, J., Mitchell, A., Nuka, G., Pesseat, S., Quinn, A. F., Sangrador-Vegas, A., Scheremetjew, M., Yong, S.-Y., Lopez, R., & Hunter, S. (2014). InterProScan 5C: Genome-scale protein function classification. Bioinformatics, 30(9), 1236–1240. 10.1093/bioinformatics/btu031

Kanazawa, T., Morinaka, H., Ebine, K., Shimada, T.L., Ishida, S., Minamino, N., Yamaguchi, K., Shigenobu, S., Kohchi, T., Nakano, A., et al. (2020). The liverwort oil body is formed by redirection of the secretory pathway. Nat Commun 11, 6152. 10.1038/s41467-020-19978-1.

Knosp, S., Kriegshauser, L., Tatsumi, K., Malherbe, L., Erhardt, M., Wiedemann, G., Bakan, B., Kohchi, T., Reski, R., & Renault, H. (2024). An ancient role for CYP73 monooxygenases in phenylpropanoid biosynthesis and embryophyte development. The EMBO Journal, 43(18), 4092–4109. 10.1038/s44318-024-00181-7

Kopylova, E., Noé, L., & Touzet, H. (2012). SortMeRNAC: Fast and accurate filtering of ribosomal RNAs in metatranscriptomic data. Bioinformatics, 28(24), 3211–3217. 10.1093/bioinformatics/bts611

Kovaka, S., Zimin, A. V., Pertea, G. M., Razaghi, R., Salzberg, S. L., & Pertea, M. (2019). Transcriptome assembly from long-read RNA-seq alignments with StringTie2. Genome Biology, 20(1), 278. 10.1186/s13059-019-1910-1

Kumar, S., Kempinski, C., Zhuang, X., Norris, A., Mafu, S., Zi, J., Bell, S. A., Nybo, S. E., Kinison, S. E., Jiang, Z., Goklany, S., Linscott, K. B., Chen, X., Jia, Q., Brown, S. D., Bowman, J. L., Babbitt, P. C., Peters, R. J., Chen, F., & Chappell, J. (2016). Molecular Diversity of Terpene Synthases in the Liverwort Marchantia polymorpha. The Plant Cell, tpc.00062.2016. 10.1105/tpc.16.00062

Lang, D., Ullrich, K. K., Murat, F., Fuchs, J., Jenkins, J., Haas, F. B., Piednoel, M., Gundlach, H., Van Bel, M., Meyberg, R., Vives, C., Morata, J., Symeonidi, A., Hiss, M., Muchero, W., Kamisugi, Y., Saleh, O., Blanc, G., Decker, E. L., … Rensing, S. A. (2018). The *Physcomitrella patens* chromosome-scale assembly reveals moss genome structure and evolution. The Plant Journal, 93(3), 515–533. 10.1111/tpj.13801

Li, F.-W., Nishiyama, T., Waller, M., Frangedakis, E., Keller, J., Li, Z., Fernandez-Pozo, N., Barker, M. S., Bennett, T., Blázquez, M. A., Cheng, S., Cuming, A. C., de Vries, J., de Vries, S., Delaux, P.-M., Diop, I. S., Harrison, C. J., Hauser, D., Hernández-García, J., … Szövényi, P. (2020). Anthoceros genomes illuminate the origin of land plants and the unique biology of hornworts. Nature Plants, 6(3), 259–272. 10.1038/s41477-020-0618-2

Li, H., Handsaker, B., Wysoker, A., Fennell, T., Ruan, J., Homer, N., Marth, G., Abecasis, G., Durbin, R., & 1000 Genome Project Data Processing Subgroup. (2009). The Sequence Alignment/Map format and SAMtools. Bioinformatics, 25(16), 2078–2079. 10.1093/bioinformatics/btp352

Love, M. I., Soneson, C., Hickey, P. F., Johnson, L. K., Pierce, N. T., Shepherd, L., Morgan, M., & Patro, R. (2020). Tximeta: Reference sequence checksums for provenance identification in RNA-seq. PLoS Computational Biology, 16(2), e1007664. 10.1371/journal.pcbi.1007664

Lundquist, P. K., Davis, J. I., & Van Wijk, K. J. (2012). ABC1K atypical kinases in plantsC: Filling the organellar kinase void. Trends in Plant Science, 17(9), 546–555. 10.1016/j.tplants.2012.05.010

Ma, J., Wang, S., Zhu, X., Sun, G., Chang, G., Li, L., Hu, X., Zhang, S., Zhou, Y., Song, C.-P., et al. (2022). Major episodes of horizontal gene transfer drove the evolution of land plants. Molecular Plant 15, 857–871. 10.1016/j.molp.2022.02.001.

Marowa, P., Ding, A., & Kong, Y. (2016). ExpansinsC: Roles in plant growth and potential applications in crop improvement. Plant Cell Reports, 35(5), 949–965. 10.1007/s00299-016-1948-4

Mata-Pérez, C., & Spoel, S. H. (2019). Thioredoxin-mediated redox signalling in plant immunity. Plant Science, 279, 27–33. 10.1016/j.plantsci.2018.05.001

Matsui, H., Iwakawa, H., Hyon, G.-S., Yotsui, I., Katou, S., Monte, I., Nishihama, R., Franzen, R., Solano, R., & Nakagami, H. (2020). Isolation of Natural Fungal Pathogens from Marchantia polymorpha Reveals Antagonism between Salicylic Acid and Jasmonate during Liverwort–Fungus Interactions. Plant and Cell Physiology, 61(2), 265–275. 10.1093/pcp/pcz187

Monte, I., Kneeshaw, S., Franco-Zorrilla, J. M., Chini, A., Zamarreño, A. M., García-Mina, J. M., & Solano, R. (2020). An Ancient COI1-Independent Function for Reactive Electrophilic Oxylipins in Thermotolerance. Current Biology, 30(6), 962–971.e3. 10.1016/j.cub.2020.01.023

Morita-Yamamuro, C., Tsutsui, T., Sato, M., Yoshioka, H., Tamaoki, M., Ogawa, D., Matsuura, H., Yoshihara, T., Ikeda, A., Uyeda, I., & Yamaguchi, J. (2005). The Arabidopsis Gene CAD1 Controls Programmed Cell Death in the Plant Immune System and Encodes a Protein Containing a MACPF Domain. Plant And Cell Physiology, 46(6), 902–912. 10.1093/pcp/pci095

Morgan M, Obenchain V, Hester J, Pagès H (2024). SummarizedExperiment: A container (S4 class) for matrix-like assays. doi:10.18129/B9.bioc.SummarizedExperiment, R package version 1.37.0, https://bioconductor.org/packages/SummarizedExperiment.

Morris, J. L., Puttick, M. N., Clark, J. W., Edwards, D., Kenrick, P., Pressel, S., Wellman, C. H., Yang, Z., Schneider, H., & Donoghue, P. C. J. (2018). The timescale of early land plant evolution. Proceedings Of The National Academy Of Sciences, 115(10). 10.1073/pnas.1719588115

Nelson, J. M., Hauser, D. A., Hinson, R., & Shaw, A. J. (2018). A novel experimental system using the liverwort *Marchantia polymorpha* and its fungal endophytes reveals diverse and context-dependent effects. New Phytologist, 218(3), 1217–1232. 10.1111/nph.15012

Ngou, B. P. M., Ding, P., & Jones, J. D. G. (2022). Thirty years of resistance: Zig-zag through the plant immune system. The Plant Cell, 34(5), 1447–1478. 10.1093/plcell/koac041

One Thousand Plant Transcriptomes Initiative. (2019). One thousand plant transcriptomes and the phylogenomics of green plants. Nature, 574(7780), 679–685. 10.1038/s41586-019-1693-2

Patro, R., Duggal, G., Love, M. I., Irizarry, R. A., & Kingsford, C. (2017). Salmon provides fast and bias-aware quantification of transcript expression. Nature Methods, 14(4), 417–419. 10.1038/nmeth.4197

Pertea, G., & Pertea, M. (2020). GFF UtilitiesC: GffRead and GffCompare. F1000Research, 9, 304. 10.12688/f1000research.23297.2

Ponce de León, I. (2023). Evolution of immunity networks across embryophytes. Current Opinion in Plant Biology, 102450. 10.1016/j.pbi.2023.102450.

Price, M. N., Dehal, P. S., & Arkin, A. P. (2010). FastTree 2 – Approximately Maximum-Likelihood Trees for Large Alignments. PLoS ONE, 5(3), e9490. 10.1371/journal.pone.0009490

Proust, H., Honkanen, S., Jones, V. A., Morieri, G., Prescott, H., Kelly, S., Ishizaki, K., Kohchi, T., & Dolan, L. (2015). RSL Class I Genes Controlled the Development of Epidermal Structures in the Common Ancestor of Land Plants. Current Biology, 26(1), 93–99. 10.1016/j.cub.2015.11.042

Quinlan, A. R. (2014). BEDToolsC: The Swiss-Army Tool for Genome Feature Analysis. Current Protocols in Bioinformatics, 47(1). 10.1002/0471250953.bi1112s47

Radhakrishnan, G. V., Keller, J., Rich, M. K., Vernié, T., Mbadinga Mbadinga, D. L., Vigneron, N., Cottret, L., Clemente, H. S., Libourel, C., Cheema, J., Linde, A.-M., Eklund, D. M., Cheng, S., Wong, G. K. S., Lagercrantz, U., Li, F.-W., Oldroyd, G. E. D., & Delaux, P.-M. (2020). An ancestral signalling pathway is conserved in intracellular symbioses-forming plant lineages. Nature Plants, 6(3), 280–289. 10.1038/s41477-020-0613-7

Redkar, A., Gimenez Ibanez, S., Sabale, M., Zechmann, B., Solano, R., and Di Pietro, A. (2022). Marchantia polymorpha model reveals conserved infection mechanisms in the vascular wilt fungal pathogen Fusarium oxysporum. New Phytologist 234, 227–241. 10.1111/nph.17909.

Rensing, S.A., Lang, D., Zimmer, A.D., Terry, A., Salamov, A., Shapiro, H., Nishiyama, T., Perroud, P.-F., Lindquist, E.A., Kamisugi, Y., et al. (2008). The Physcomitrella Genome Reveals Evolutionary Insights into the Conquest of Land by Plants. Science 319, 64–69. 10.1126/science.1150646.

Rensing, S. A. (2018). Great moments in evolution: the conquest of land by plants. Current Opinion In Plant Biology, 42, 49–54. 10.1016/j.pbi.2018.02.006

Rich, M. K., Vigneron, N., Libourel, C., Keller, J., Xue, L., Hajheidari, M., Radhakrishnan, G. V., Le Ru, A., Diop, S. I., Potente, G., Conti, E., Duijsings, D., Batut, A., Le Faouder, P., Kodama, K., Kyozuka, J., Sallet, E., Bécard, G., Rodriguez-Franco, M., … Delaux, P.-M. (2021). Lipid exchanges drove the evolution of mutualism during plant terrestrialization. Science, 372(6544), 864–868. 10.1126/science.abg0929

Robinson, M. D., McCarthy, D. J., & Smyth, G. K. (2010). edgeRC: A Bioconductor package for differential expression analysis of digital gene expression data. Bioinformatics, 26(1), 139–140. 10.1093/bioinformatics/btp616

Robinson, M. D., & Oshlack, A. (2010). A scaling normalization method for differential expression analysis of RNA-seq data. Genome Biology, 11(3), R25. 10.1186/gb-2010-11-3-r25

Romani, F., Banić, E., Florent, S.N., Kanazawa, T., Goodger, J.Q.D., Mentink, R.A., Dierschke, T., Zachgo, S., Ueda, T., Bowman, J.L., et al. (2020). Oil Body Formation in Marchantia polymorpha Is Controlled by MpC1HDZ and Serves as a 109 Defense against Arthropod Herbivores. Current Biology 30, 2815–2828.e8. 10.1016/j.cub.2020.05.081.

Romani, F., Flores, J.R., Tolopka, J.I., Suárez, G., He, X., and Moreno, J.E. (2022). Liverwort oil bodies: diversity, biochemistry, and molecular cell biology of the earli est secretory structure of land plants. Journal of Experimental Botany 73, 4427 4439. 10.1093/jxb/erac134.

Ros-Moner, E., Jiménez-Góngora, T., Villar-Martín, L., Vogrinec, L., González-Miguel, V. M., Kutnjak, D., & Rubio-Somoza, I. (2024). Conservation of molecular responses upon viral infection in the non-vascular plant Marchantia polymorpha. Nature Communications, 15(1). 10.1038/s41467-024-52610-0

Rosado, C. J., Kondos, S., Bull, T. E., Kuiper, M. J., Law, R. H. P., Buckle, A. M., Voskoboinik, I., Bird, P. I., Trapani, J. A., Whisstock, J. C., & Dunstone, M. A. (2008). The MACPF/CDC family of pore-forming toxins. Cellular Microbiology, 10(9), 1765–1774. 10.1111/j.1462-5822.2008.01191.x

Schornack, S., & Kamoun, S. (2023). EVO-MPMI: From fundamental science to practical applications. Current Opinion In Plant Biology, 76, 102469. 10.1016/j.pbi.2023.102469

Shumate, A., & Salzberg, S. L. (2021). LiftoffC: Accurate mapping of gene annotations. Bioinformatics, 37(12), 1639–1643. 10.1093/bioinformatics/btaa1016

Stanke, M., Schöffmann, O., Morgenstern, B., & Waack, S. (2006). Gene prediction in eukaryotes with a generalized hidden Markov model that uses hints from external sources. BMC Bioinformatics, 7(1), 62. 10.1186/1471-2105-7-62

Stanke, M., Diekhans, M., Baertsch, R., & Haussler, D. (2008). Using native and syntenically mapped cDNA alignments to improve *de novo* gene finding. Bioinformatics, 24(5), 637–644. 10.1093/bioinformatics/btn013

Suire, C., Bouvier, F., Backhaus, R.A., Bégu, D., Bonneu, M., and Camara, B. (2000). Cellular Localization of Isoprenoid Biosynthetic Enzymes in Marchantia polymorpha Uncovering a New Role of Oil Bodies. Plant Physiology 124, 971 978. 10.1104/pp.124.3.971.

Wang, L., Wan, M.-C., Liao, R.-Y., Xu, J., Xu, Z.-G., Xue, H.-C., Mai, Y.-X., & Wang, J.-W. (2023). The maturation and aging trajectory of Marchantia polymorpha at single-cell resolution. Developmental Cell, 58(15), 1429–1444.e6. 10.1016/j.devcel.2023.05.014

Wang, Y., Tang, H., DeBarry, J. D., Tan, X., Li, J., Wang, X., Lee, T. -h., Jin, H., Marler, B., Guo, H., Kissinger, J. C., & Paterson, A. H. (2012). MCScanXC: A toolkit for detection and evolutionary analysis of gene synteny and collinearity. Nucleic Acids Research, 40(7), e49–e49. 10.1093/nar/gkr1293

Yokota, T., Ohnishi, T., Shibata, K., Asahina, M., Nomura, T., Fujita, T., Ishizaki, K., & Kohchi, T. (2017). Occurrence of brassinosteroids in non-flowering land plants, liverwort, moss, lycophyte and fern. Phytochemistry, 136, 46–55. 10.1016/j.phytochem.2016.12.020

Yotsui, I., Matsui, H., Miyauchi, S., Iwakawa, H., Melkonian, K., Schlüter, T., Michavila, S., Kanazawa, T., Nomura, Y., Stolze, S. C., Jeon, H., Yan, Y., Harzen, A., Sugano, S. S., Shirakawa, M., Nishihama, R., Ichihashi, Y., Ibanez, S. G., Shirasu, K., … Nakagami, H. (2023). LysM-mediated signaling in Marchantia polymorpha highlights the conservation of pattern-triggered immunity in land plants. Current Biology, 33(17), 3732–3746.e8. 10.1016/j.cub.2023.07.068

Zhang, J., Fu, X.-X., Li, R.-Q., Zhao, X., Liu, Y., Li, M.-H., Zwaenepoel, A., Ma, H., Goffinet, B., Guan, Y.-L., et al. (2020). The hornwort genome and early land plant evolution. Nat. Plants 6, 107–118. 10.1038/s41477-019-0588-4.

Zhou, X., & Stephens, M. (2012). Genome-wide efficient mixed-model analysis for association studies. Nature Genetics, 44(7), 821–824. 10.1038/ng.2310

