## Supplementary material for "Horizontal gene transfers and terpene metabolism drive plant-fungal interaction in *Marchantia polymorpha*": legends_sup_data.docx

**Data S1:** Detailed table of the Hierarchical clustering of Tak-1’s significantly differentially expressed genes during *C. nymphaeae* infection (adjusted p ≤ 0.05; log2 fold change [logFC] ≥ 1) at 0 and 3 days post inoculation (dpi) (Figure 3.a). Variance stabilized row-centered counts are shown. Additional information on these genes is specified: gene symbol, functional annotation, pangenomic compartment of the genes, differential expression during the infection with *P. palmivora* (adjusted p-value ≤ 0.05; absolute log2 fold change [logFC] ≥ 1), and differential expression under different single or cumulated abiotics stresses (N=nitrogen deficiency, S=salt, L=light, M=mannitol, C=cold, D=dark, H=heat).

**Data S2:** Full table of the functional enrichments (GO and IPR terms) on Tak-1 up and down regulated genes in response to *C. nymphaeae* at 3 dpi (enrichment cutoff of 0.01). The IPR enrichment were then curated to remove the redundant terms in order to produce the Figure 3.b.

**Data S3: Cross referencing of the single copy** orthogroups differentially expressed in both *M. polymorpha* and *N. benthamiana* in response to *P. palmivora* (Carella et al. 2019) with *M. polymorpha* differentially expressed genes in response to *C. nymphaeae*, and functional enrichment on the genes commonly up regulated in *M. polymorpha* in response to both pathogens and in *N. benthamiana* in response to *P. palmivora*.

**Data S4:** Table of the differential expression of all *M. polymorpha* genes in Tak-1 and CA at 3 and 6 dpi with *C. nymphaeae*. Information about the GWAS candidate genes, the genome environment association candidate genes and pangenomic compartment (Beaulieu et al. 2024), the cell cluster assignation by single cell RNAseq (Wang et al. 2023), the differential expression under P. palmivora infection (Carella et al. 2019) and under various abiotic stresses (Wen Tan et al. 2023), as well as the functional annotations are specified. Correspondance between CA and Tak-1 genes can lead to multiple lines for a single gene in one accession (but multiple corresponding genes in the other accession).

**Data S5:** Detailed table of the Hierarchical clustering of CA’s significantly differentially expressed genes during *C. nymphaeae* infection (adjusted p ≤ 0.05; log2 fold change [logFC] ≥ 1) at 0, 3 and 6 days post inoculation (dpi) (Figure 3.c). Variance stabilized row-centered counts are shown. Additional information on these genes is specified: gene symbol, functional annotation, pangenomic compartment of the genes, differential expression during the infection with *P. palmivora* (adjusted p-value ≤ 0.05; absolute log2 fold change [logFC] ≥ 1), and differential expression under different single or cumulated abiotics stresses (N=nitrogen deficiency, S=salt, L=light, M=mannitol, C=cold, D=dark, H=heat).

**Data S6:** Full table of the functional enrichments (GO and IPR terms) on CA up regulated genes at 3 and 6 dpi in response to *C. nymphaeae* (enrichment cutoff of 0.01). Not enough genes were down regulated to perform an enrichment on them. The IPR enrichment were then curated to remove the redundant terms in order to produce the Figure 3.d.

**Data S7:** List of genomic regions and genes associated with the four phenotypes considered at 6dpi: thallus area of *C. nymphaeae* inoculated plants, thallus area of mock inoculated plants, symptoms area (brown) and ratio of symptoms area on thallus area (brown percentage). The genome wide association study was performed on 77 *M. polymorpha* ssp. *ruderalis* accessions.

**Data S8**: Macros used for *M. polymorpha* phenotyping.

**Data S9:** Species contained in a database of 261 genomes from all the main clades of land plants, and used for phylogenetic analyses.

**Figure S1:** Phenotypic variability in response to *C. nymphaeae* infection, within the *M. polymorpha* collection.	*C. nymphaeae* infection causes maceration of the thallus resulting in browning. In the most susceptible accessions, white hyphae are visible on the surface of the thallus. Nor-E : Norwich-E, Tou-C : Toulon-C, Gil-1: Gilesgate-1, Voe-B : Voewood-B, Cam-2 : Cambridge-2, Tak-1: Takaragaike-1, Bul-B: Bulan-B.

**Figure S2:** Phylogeny of *M. polymorpha*’s GH88 gene (Mp3g19320). This tree was computed with FastTree.

**Table S1** : Primers used to identify *Colletrotrichum spp*. (based on Damm et al., 2014).
