## Supplementary material for "Horizontal gene transfers and terpene metabolism drive plant-fungal interaction in *Marchantia polymorpha*": Sup_Table_S1_Primers used to identify Colletrotrichum spp.docx

| **Supplementary Table 1 : Primers used to identify *Colletrotrichum spp*. (based on Damm et al., 2014)** | | |
| --- | --- | --- |
| **Name of   targeted sequence** | **Forward (F) /Reverse (R)** | **Primer sequence** |
| **ITS-1** | F | CTTGGTCATTTAGAGGAAGTAA |
|  | R | TCCTCCGCTTATTGATATGC |
| **GAPDH** | F | GCCGTCAACGACCCCTTCATTGA |
|  | R | GGGTGGAGTCGTACTTGAGCATGT |
| **CHS1 (partial)** | F | TGGGGCAAGGATGCTTGGAAGAAG |
|  | R | TGGAAGAACCATCTGTGAGAGTTG |
| **HIS3** | F | AGGTCCACTGGTGGCAAG |
|  | R | AGCTGGATGTCCTTGGACTG |
| **ACT** | F | ATGTGCAAGGCCGGTTTCGC |
|  | R | TACGAGTCCTTCTGGCCCAT |
| **TUB** | F | ACCCTCAGTGTAGTGACCCTTGGC |
|  | R | ACCCTCAGTGTAGTGACCCTTGGC |
