## Supplementary figures and images for "Horizontal gene transfers and terpene metabolism drive plant-fungal interaction in *Marchantia polymorpha*"

### Supp Figure 1.png

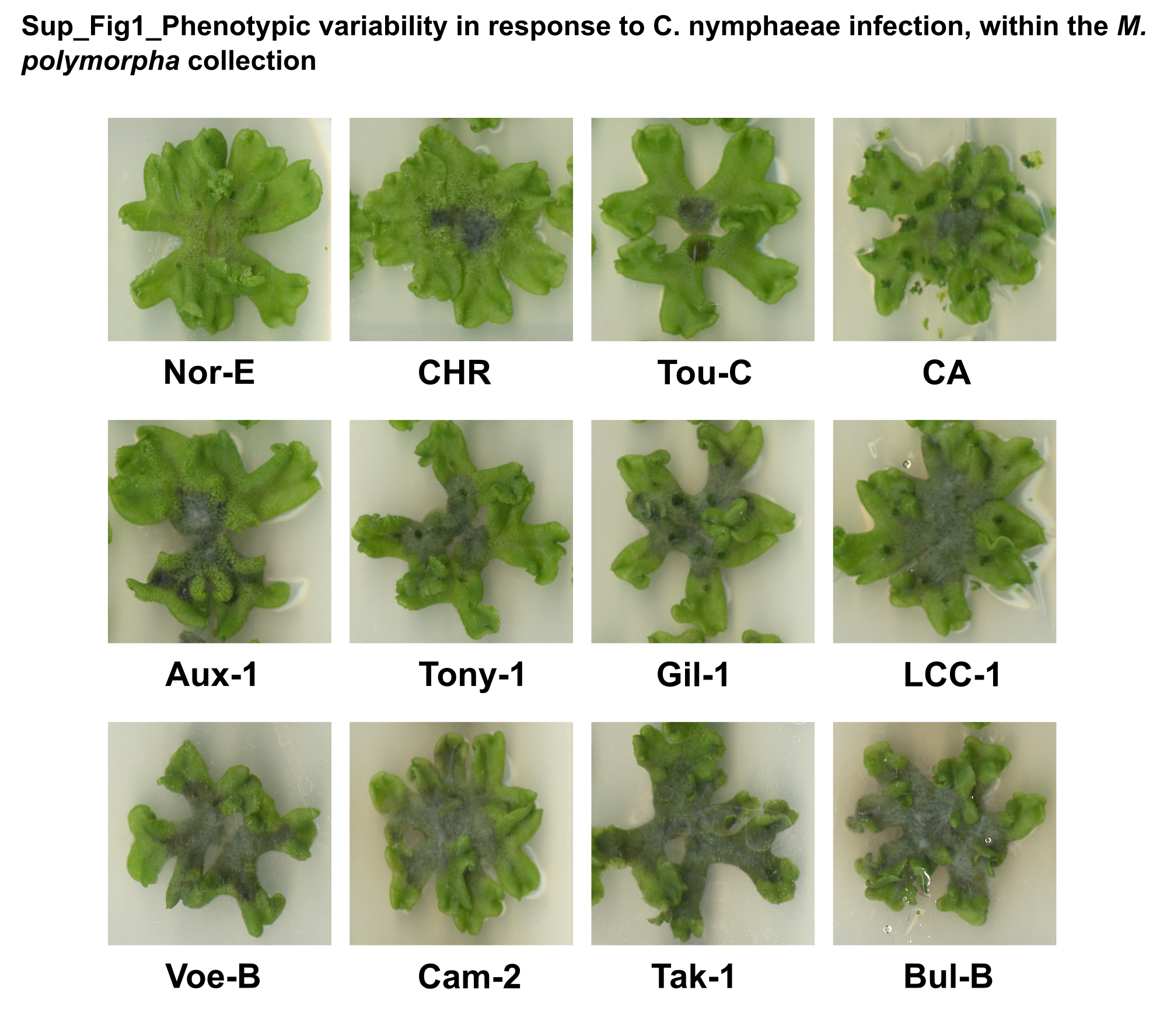

### Supp Figure 2.png

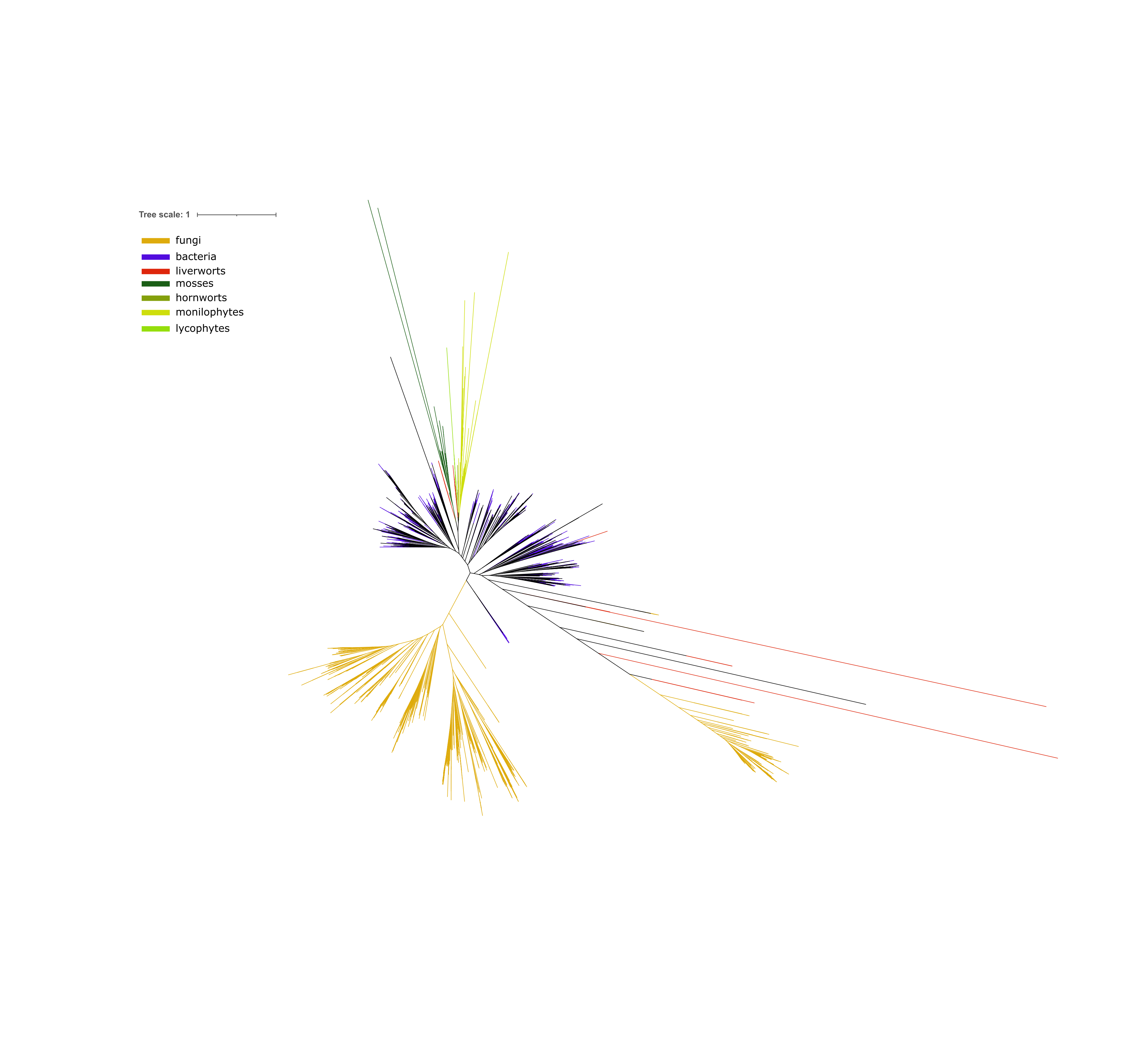
